# Simultaneous readout of multiple FRET pairs using photochromism

**DOI:** 10.1101/2021.01.06.425528

**Authors:** Thijs Roebroek, Wim Vandenberg, François Sipieter, Siewert Hugelier, Christophe Stove, Jin Zhang, Peter Dedecker

## Abstract

Förster resonant energy transfer (FRET) is a powerful mechanism to probe associations *in situ*. Simultaneously performing more than one FRET measurement can be challenging due to the spectral band-width required for the donor and acceptor fluorophores. We present an approach to distinguish overlapping FRET pairs based on the photochromism of the donor fluorophores, even if the involved fluorophores display essentially identical absorption and emission spectra. We develop the theory underlying this method and validate our approach using numerical simulations. To apply our system, we develop rsAKARev, a photochromic biosensor for cAMP-dependent kinase (PKA), and combine it with the spectrally-identical biosensor EKARev, a reporter for ERK kinase activity, to deliver simultaneous readout of both activities in the same cell. We further perform multiplexed PKA, ERK, and calcium measurements by including a third, spectrally-shifted biosensor. Our work demonstrates that exploiting donor photochromism in FRET can be a powerful approach to simultaneously read out multiple associations within living cells.

## Introduction

Förster resonance energy transfer (FRET) is a powerful mechanism to probe associations *in situ*. FRET experiments are performed by labeling the system of interest with donor and acceptor fluorophores, and then using the extent of the energy transfer to probe the distances between and relative orientations of both fluorophores [1]. FRET-based approaches have gained tremendous popularity for association studies, and can be performed using both genetically-encoded fluorophores and organic dyes [2]. It is also the key mechanism underlying a class of genetically-encoded biosensors that convert the presence of a particular chemical stimulus into a change in FRET [3–6].

Most FRET experiments use a fluorescent donor and acceptor, which has the advantage that the FRET efficiency can be estimated based on the ratio between the donor-excited emission from both fluorophores [7–10]. The accuracy of this estimation depends on the ease with which the donor and acceptor can be spectrally separated from one another. The choice of specific FRET pairs is usually driven by a compromise between achieving a clearly observable FRET efficiency, which requires sufficient spectral overlap, and maintaining a low level of cross-talk between channels, which is made easier when the probes are more spectrally separated. As a result, the donor and acceptor tend to occupy broad regions of the visual spectrum, complicating the combination of FRET measurements with the use of other probes. Measuring multiple FRET-based constructs is further complicated if these constructs share fluorophores emitting in the same spectral band. For example, many FRET biosensors consist of a cyan and a yellow fluorescent protein, making it more difficult to combine multiple biosensors to follow e.g. cross-talk between signaling pathways. Using more than one of these probes typically requires that separate measurements are performed for each probe, possibly in a high-throughput manner [11].

A number of strategies can facilitate FRET multiplexing [12]. The conceptually easiest approach is to select two or more FRET pairs with minimal spectral overlap, though this is limited by the available probes and the usable spectral range [13]. It may be possible to share a single fluorophore among two FRET pairs, reducing the total number of fluorophores from four to three [14]. Other approaches include the use of non-fluorescent acceptors combined with fluorescence lifetime imaging [15–17], the use of homo-FRET combined with alternative readouts such as fluorescence anisotropy [18], and the separation of spectrally overlapping pairs based on orthogonal information, such as the spatial localization of the probes within the cell [19].

Several strategies for probe multiplexing have relied on exploiting the photochemistry of the fluorophores. Photochromic labels, for example, display light-induced and reversible ‘on-off’ switching of their fluorescence emission [20, 21]. Upon excitation, these probes display a bright fluorescence emission that disappears in time, though the original fluorescence can be rapidly recovered by irradiating with light of a shorter wavelength. These dynamics can be used to separate the signal of a photochromic label from that of a non-photochromic label or from the autofluorescence, or can be used to separate two or more photochromic labels that switch with different kinetics [22–26]. However, there has been comparatively little work on the evaluation of such strategies for the simultaneous measurement of multiple FRET pairs. In contrast, photochromism of the acceptor has been used as a way to provide a more accurate FRET quantification [27, 28] or to measure multiple single-molecule donor-acceptor distances within the same complex [29]. Previous work has also investigated the use of a photochromic donor, where the FRET efficiency can be estimated by determining the rate of the photochromism [30], or to facilitate the anisotropy-based readout of homo-FRET [31]. Association-induced fluorescence dynamics have also been used to provide diffraction-unlimited imaging of biosensors [32].

In this contribution we describe the use of donor photochromism to simultaneously measure more than one FRET pair, even when the donors and acceptors display complete spectral overlap. We commence by developing the theory underlying our method, and validate its performance using numerical simulations. We then develop a photochromic biosensor for cAMP-dependent protein kinase (PKA), which we call rsAKARev, and use it to demonstrate our approach through the simultaneous measurement of PKA and extracellular signal-regulated kinase (ERK) activity in live cells. Finally, we demonstrate the versatility of our approach by combining these sensors with a spectrally-separated calcium biosensor to perform simultaneous biosensing of three different signaling activities in live cells.

## Theory

### Quantification of the FRET signal

The goal of a FRET measurement is to quantify the energy transfer between the donor and acceptor fluorophore. We assume that this measurement is performed using sensitized imaging, where the donor is excited and the emission from the donor (*S*_*DD*_) and acceptor (*S*_*DA*_) are measured simultaneously or in rapid succession. As a control, and to estimate the effects of photodestruction, one usually also measures the acceptor emmission uppon direct excitation (*S*_*AA*_). A full FRET acquisition therefore consists of three measurements {*S*_*DD*_,*S*_*DA*_,*S*_*AA*_} at every timepoint and at every pixel of the detector.

The *S*_*DA*_ signal is usually contaminated by donor bleed-through and direct acceptor cross-excitation. These contributions can be quantified by using explicit correction factors *α* and *δ* [9]:

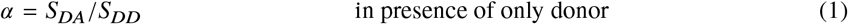

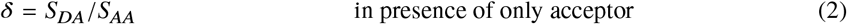

These correction factors can be determined using control measurements, or can be estimated given knowledge of the fluorophore spectra and the spectral sensitivity of the instrument. In general, direct control measurements are preferable since these will then be tailored to the specifics of the instrument. We assume that direct excitation of the acceptor does not lead to appreciable emission of the donor fluorophore in the acceptor channel.

Not all experimenters choose to determine these correction factors, instead quantifying the energy transfer using the ‘raw’ emission ratio *R*

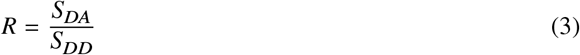

which will already allow the detection of relative changes in the FRET efficiency. If available, however, we can use the correction factors to calculate the sensitized emission *F*_*c*_, which is the donor-excited emission from the acceptor corrected for cross-talk:

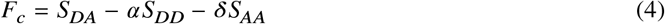

Given *F*_*c*_, the FRET response can be estimated using the sensitized emission ratio *R*_*c*_:

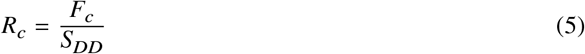

*R*_*c*_ can be compared across different instruments and is more sensitive to changes in FRET than the emission ratio *R*. However, it is not unambiguously related to the absolute FRET efficiency (Θ) since it fails to take into account the different brightnesses of the donor and acceptor fluorophores and/or the different collection efficiencies of the instrument. The introduction of an additional correction factor *γ* [9] makes it possible to calculate the FRET efficiency using

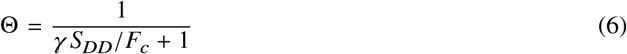

as is derived in section 1 of the supplementary information. *γ* can be estimated from model spectra, or more reliably from additional control experiments [10].

For some systems, such as genetically-encoded biosensors, changes in the observed FRET efficiency arise from the interconversion of the system between a low- and a high-FRET state, rather than from a continuous variation of the FRET efficiency. For this reason we will refer to the result of Equation (6) as the ‘apparent FRET efficiency’ when discussing measurements on FRET biosensors.

### Introducing a photochromic donor

The core idea of our method is that two different FRET pairs can be distinguished based on the photochromism of the donors, conceptually shown in Figure 1a. We therefore extend our model to the case where the donor is a photochromic fluorophore. To probe this photochromism, we expand our measurement scheme with two irradiation periods that serve to switch the donor fluorophore between fluorescent and non-fluorescent states (Figure 1b). This consists of a short-wavelength light pulse and fluorescence acquisition to capture the fluorescent state, followed by a longer-wavelength light pulse and another fluorescence acquisition to capture the extent of the off-switching. The net result is that every FRET acquisition is now replaced with two acquisitions, performed at the end of each irradiation period, for a total of six measurements for each FRET timepoint (two measurements each for *S*_*DD*_, *S*_*DA*_, and *S*_*AA*_). *S*_*AA*_ usually varies slowly because it is affected only by photodestruction or large structural changes in the sample, and can therefore be measured at a reduced frequency.

**Figure 1:**
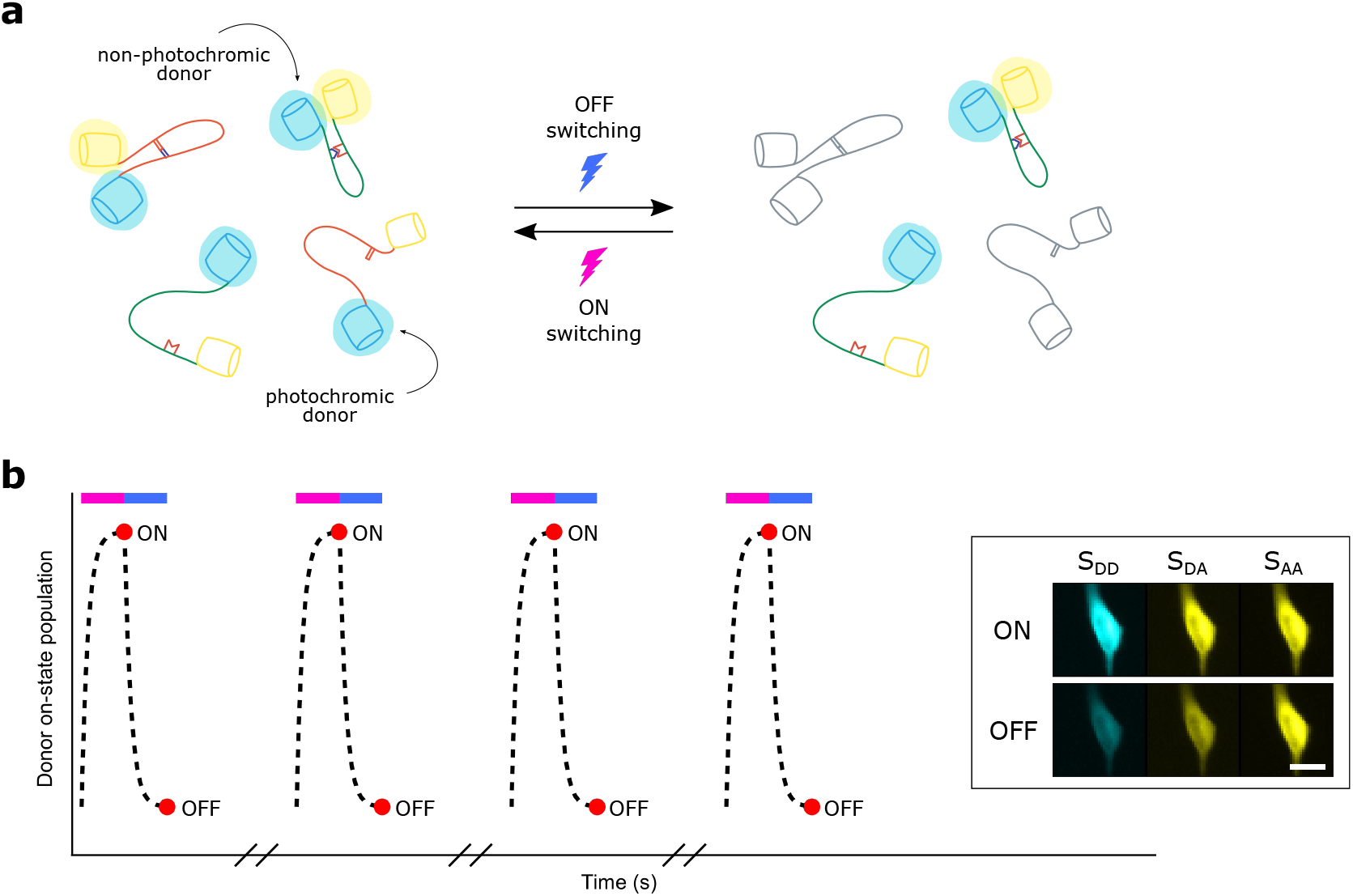
(a) Overview of the concept: two FRET pairs can be distinguished by probing the photochromism of the donors, since only photochromic donors react to the off- and on-switching light. (b) Example irradiation scheme for a cyan photochromic protein, showing the donor on-state population (dotted line) during violet and blue light irradiation (colored bars). The red dots show the fluorescence acquisitions that are performed. The inset shows representative fluorescence images of a HeLa cell, acquired in all three measurement channels. Scale bar, 20 μm. In practice, the light-induced off-switching does not need to be as complete as shown here.

Off-switching of the donor fluorophore results in a decrease in *S*_*DD*_, *S*_*DA*_ and *F*_*c*_. We quantify this using the photoswitching ratios *ρ*_*D*_ and *ρ*_*A*_, given by

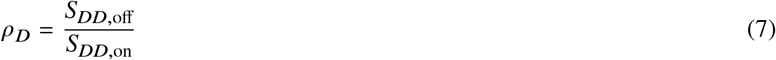

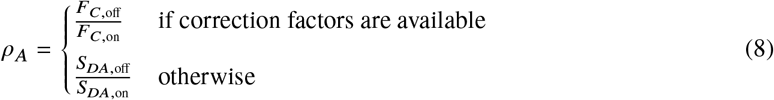

where the ‘on’ and ‘off’ acquisitions are those shown in Figure 1b. We provide two alternative definitions of *ρ*_*A*_ in Equation (8) depending on whether correction factors for cross-talk are available. Our methodology remains valid for both approaches as long as these definitions are used consistently.

The photoswitching ratios *ρ*_*D*_ and *ρ*_*A*_ are thus the fractions of the original fluorescence that are left after the off-switching irradiation has been applied, with smaller values implying more pronounced off-switching. They depend on the specifics of the donor fluorophore and on the total light dose that is delivered, but also on the FRET efficiency since this process competes with the photochromism. This aspect was leveraged in the development of psFRET [30], and likewise must be taken into account when separating two FRET pairs based on photochromism.

In the general case, the FRET-dependence of the photoswitching ratios *ρ*_*D*_ and *ρ*_*A*_ must be determined experimentally by measuring these for varying (sensitized) emission ratios or FRET efficiencies. This can be done simply by performing an experiment where the FRET pair is expressed and *ρ*_*D*_ and *ρ*_*A*_ are followed as a function of emission ratio or FRET efficiency. This approach will also work in the more general case where the emission contrast arises not just due to energy transfer, but also through other contributions such as a change in brightness of the donor.

This process can be simplified when suitable assumptions can be made about the system. For a simple FRET process with negligible or perfectly corrected cross-talk in which only a single species is present,

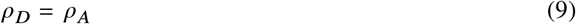

since the acceptor emission is proportional to the amount of the donor in the fluorescent state. If we further assume that the off-switching can be described by an intrinsic quantum yield then the photoswitching ratio is given by

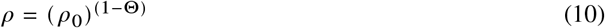

as is derived in section 2 of the supplementary information. Equation (10) shows that the full dependence is known if one simply measures *ρ*_0_, the photoswitching ratio in the absence of FRET, using the same settings for the instrument. As we show in the supplementary information, this consideration can be expanded to photochromic molecules that do not display complete off-switching but instead reach an equilibrium or plateau level.

While these results can simplify the characterization of the FRET pairs, our method will still work if these assumptions cannot be made. Residual cross-talk of direct acceptor excitation into the *S*_*DA*_ channel, for example, will increase *ρ*_*A*_ since this contribution shows no fluorescence modulation. The derived models also do not apply to FRET-based biosensors, which exist as two interconverting species (active and inactive forms of the sensor). If no simplifying assumptions can be made, the dependence of *ρ*_*A*_ and *ρ*_*D*_ on the (sensitized) emission ratio or FRET efficiency will need to be determined over the full range of values, for each FRET pair and off-switching light dose used in the experiment.

### Separating photochromic FRET pairs

We now assume that the sample is labeled with two FRET pairs, where the absorption and emission spectra are highly similar yet the donors differ in their photochromism. This situation can also arise when two different interactions are probed using two donors and a single shared acceptor. The recorded fluorescence signals are simply the sum of the signals from the individual components

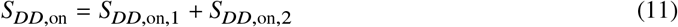

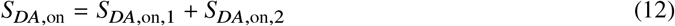

where ‘1’ and ‘2’ indicate the first and second FRET pair. After irradiation with off-switching light, we also obtain

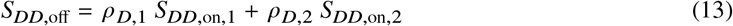

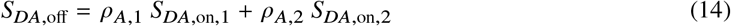

Our measurement provides the *S*_*DD*_ and *S*_*DA*_ signals before and after off-switching. If we assume that all four photoswitching ratios *ρ*_*A*_ and *ρ*_*D*_ are known, then these equations constitute a system of four equations and four unknowns, which can be solved to obtain *S*_*DD*,on_ and *S*_*DA*,on_ for both species. These values can then be readily used to obtain the (sensitized) emission ratios for each pair using Equations (3) or (5), or the FRET efficiency using Equation (6). In principle, the same approach could also be used to separate two FRET pairs consisting of a single photochromic donor and a fluorescent and a dark (non-fluorescent) acceptor.

In practice, we do not know the four *ρ*_*D*_ and *ρ*_*A*_ values as these depend on the sought-after (sensitized) emission ratio or FRET efficiency. However, the previous section considered how to determine the relationship between these parameters. Given that information, we recommend the use of an iterative procedure, in which the (sensitized) emission ratio or FRET efficiency is calculated starting from initial guesses for the photoswitching ratios. This value is then used to obtain better guesses for the photoswitching ratios, and this procedure is repeated until these values converge, which occurs rapidly in practice. A more detailed discussion of the algorithm used in this work can be found in section 3 of the supplementary information.

We devised a series of numerical simulations to test the validity of our approach. We assumed a sample labeled with two spectrally indistinguishable FRET pairs, where one of the donors is photochromic and can be described using Equation (10), and the other does not show appreciable photochromism. We further assumed different brightness levels, reflected in the number of photons that would be acquired in the ‘on’ acquisition if no FRET were to occur. Noise was included in our measurement by simulating the Poisson noise intrinsic to the photon detection process. For each condition, we ran 5,000 simulations where we selected random ground-truth FRET efficiencies between 0 and 0.6 for each FRET pair, and generated two simulated fluorescence acquisitions, before and after off-switching (cfr. the measurement scheme shown in Figure 1b). This data was then analyzed using our methodology, resulting in estimates for the FRET efficiency of each of the pairs. The accuracy of these results was then quantified using the mean absolute deviation, given by

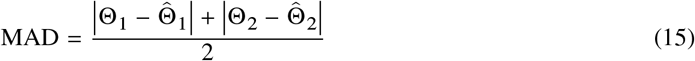

where Θ_*i*_ are the ground truth FRET efficiencies and 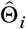 are the FRET efficiencies estimated by our method. As a negative control we also included the result of an analysis that simply generated random FRET efficiencies.

Our simulations show that our analysis works well already at low light levels, providing accurate results when 1,000 or more photons are detected, and less accurate but useful results already at signal levels as low as 250 photons (Figure 2a). We also investigated to what extent the difference in photochromism performance of the FRET donors (Δ*ρ*) determines the accuracy of the analysis (Figure 2b and Supplementary Figure 1). We find that this depends on the expected FRET efficiency and the donor photoswitching ratio in the abscence of FRET (*ρ*_0_). However, a *ρ*_0_ of 0.3 or lower, readily achievable with e.g. photochromic fluorescent proteins, works well over a broad range of FRET efficiencies. Furthermore, *ρ*_0_ can be increased or decreased by adjusting the duration or intensity of the off-switching illumination, allowing the conditions to be tailored to the measurement. Finally, we note that further increases in accuracy are possible by including more fluorescence acquisitions as part of the off-switching (adding additional fluorescence acquisitions to each of the measurement cycles shown in Figure 1b).

**Figure 2:**
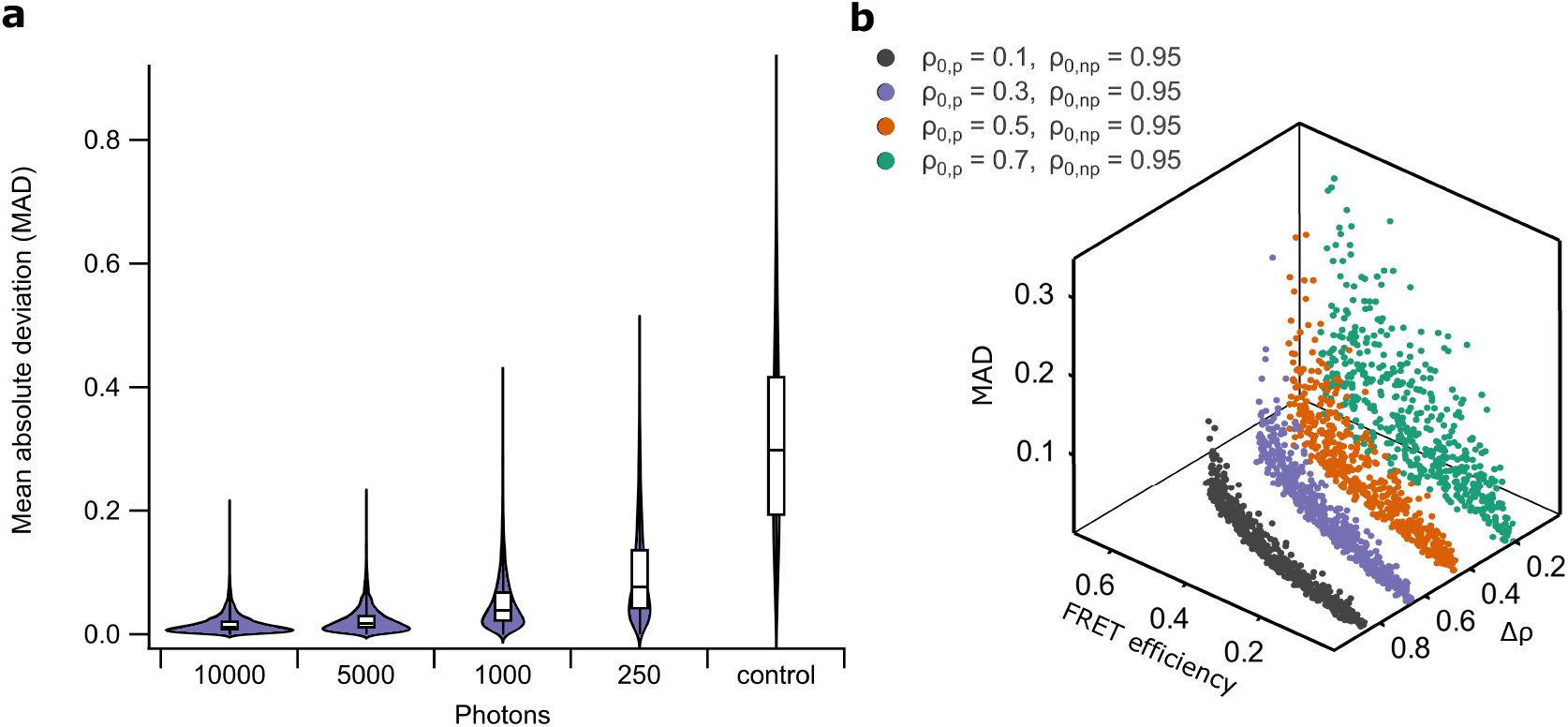
Performance of our method on simulated data. (a) Combined violin and box-plot of the mean absolute deviation (MAD) of our analysis approach for different numbers of detected photons. Data was generated by performing 5,000 independent simulations for each category. (b) Scatter plot showing the analysis performance for different FRET efficiencies and photoswitching ratios (in the absence of FRET) of the photochromic (*ρ*_0,*p*_) and non-photochromic (*ρ*_0,*np*_) donor. 2D projections of this data are shown in Supplementary Figure 1.

In principle, our methodology can be used to separate more than two overlapping FRET pairs, by adding additional terms to Equations (11) to (14). Each additional pair will then require one additional fluorescence acquisition per switching cycle.

## Results and Discussion

### rsAKARev: a photochromic biosensor for PKA

We decided to experimentally validate our method by developing a photochromic FRET-based biosensor for cAMP-dependent protein kinase (PKA) activity, called rsAKARev (reversibly switchable A-kinase activity reporter with EV-linker). We obtained this biosensor by replacing the donor and acceptor in AKAR3ev [33] with the cyan fluorescent protein mTFP0.7 and the yellow fluorescent protein cpVenus172. mTFP0.7 is a variant of the bright and photostable mTFP1 [34] that shows efficient photochromism with little fatigue [35]. cpVenus172 is a circular permutant of Venus that has been reported to result in an improved FRET contrast [36–38]. To evaluate our unmixing strategy, we chose to combine rsAKARev with the non-photochromic biosensor EKARev [33], which reports on extracellular signal-regulated kinase (ERK) activity and consists of the cyan donor ECFP and yellow acceptor YPet. Our choice of sensors reflects the fundamental interest in the interactions between the conserved cAMP/PKA and MAPK/ERK signaling pathways, which regulate various physiological processes including cell proliferation and growth [39, 40].

To characterize the performance of our biosensors, we performed measurements on cells expressing either rsAKARev or EKARev, and submitted them to the measurement scheme shown in Figure 1b. As expected, we found that the fluorescent state of the rsAKARev donor fluorophore, mTFP0.7, readily depopulates upon irradiation with blue light and can be efficiently recovered through weak irradiation with violet light (Figure 3a). This donor photochromism leads to a large and reversible fluorescence contrast in the donor and acceptor emission with minimal loss of fluorescence intensity (Figure 3b). In contrast, the EKARev donor, ECFP, displays a much more limited modulation of the fluorescence signal (Figure 3c).

**Figure 3:**
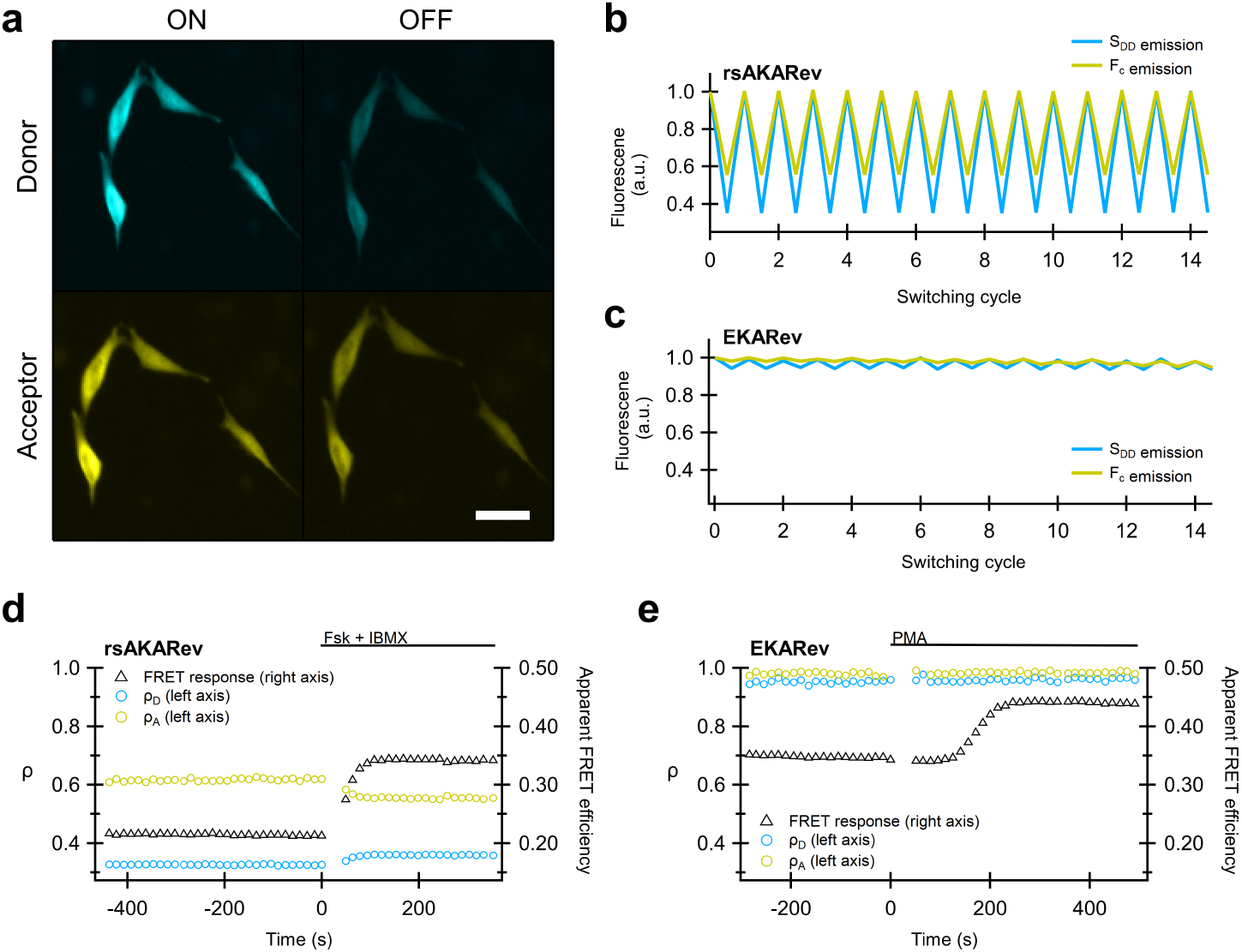
Photochromism behavior of rsAKARev and EKARev expressed in HeLa cells. (a) Representative fluorescence images of the donor (*S*_*DD*_) and acceptor (*S*_*DA*_) fluorescence of rsAKARev before and after off-switching. Scale bar, 40 μm (b,c) Time-course of the donor and sensitized acceptor emission of rsAKARev (b) and EKARev (c) during multiple irradiation cycles. (d,e) Representative responses of a HeLa cell expressing rsAKARev (d) or EKARev (e) during stimulation of PKA or ERK activity. Time-traces are colored as indicated. PKA activity was stimulated with forskolin (Fsk, 50 μM) and IBMX (100 μM), ERK activity with PMA (1 μM).

We next evaluated how the activation state of each biosensor affected its fluorescence emission. Cells were treated with pharmacological compounds to evoke either PKA or ERK kinase activity, and the FRET response of the respective biosensor was followed over time (Figure 3d,e). This response was determined via conventional FRET analysis using only the ‘on’ fluorescence acquired after every violet irradiation step in Figure 1b.

Forskolin-(Fsk) and 3-isobutyl-1-methylxanthine (IBMX)-mediated elevation of intracellular cAMP strongly increased the FRET response of rsAKARev. Likewise, stimulation of cells with phorbol-12-myristate-13-acetate (PMA), which acts on ERK via upstream kinases [41, 42], elicited a strong EKARev response. As expected, our data additionally revealed that changes in PKA activity also lead to changes in the photo-switching ratios *ρ*_*D*_ and *ρ*_*A*_ (Figure 3d). Conversely, Figure 3e shows that the irradiation-induced fluorescence dynamics of EKARev are small and largely independent of the activation state of the sensor, consistent with the photostatic nature of ECFP.

As we discussed in the theory, separating overlapping FRET pairs based on their photochromism requires knowledge of the relationship between the photoswitching ratio and the FRET efficiency. For well-behaved FRET processes, this can be done using only a single reference measurement and the analytical expression of the dependence (e.g. Equation (10)). However, systems that consist of an equilibrium between multiple discrete states, such as the active and inactive state of a biosensor, cannot be described using this relationship. Furthermore, cyan fluorescent proteins commonly show a complicated spectroscopic behavior such as multi-exponential excited state lifetime decays [43]. We therefore resorted to direct measurement of the full dependence in cells expressing either rsAKARev or EKARev (Figure 4). We found that the sought-after relationship could be empirically described using double-exponential (rsAKARev) or straight line (EKARev) fits, providing a model that was used in the downstream analysis. Figure 4b does show that the photoswitching ratio increases with decreasing apparent FRET efficiency, which is contrary to what one would expect for a two-state FRET system. This is likely due to a suboptimal value for the acceptor bleed-through correction, though this does not interfere in the separation analysis which follows since its effects are automatically taken into account via this *in situ* calibration.

**Figure 4:**
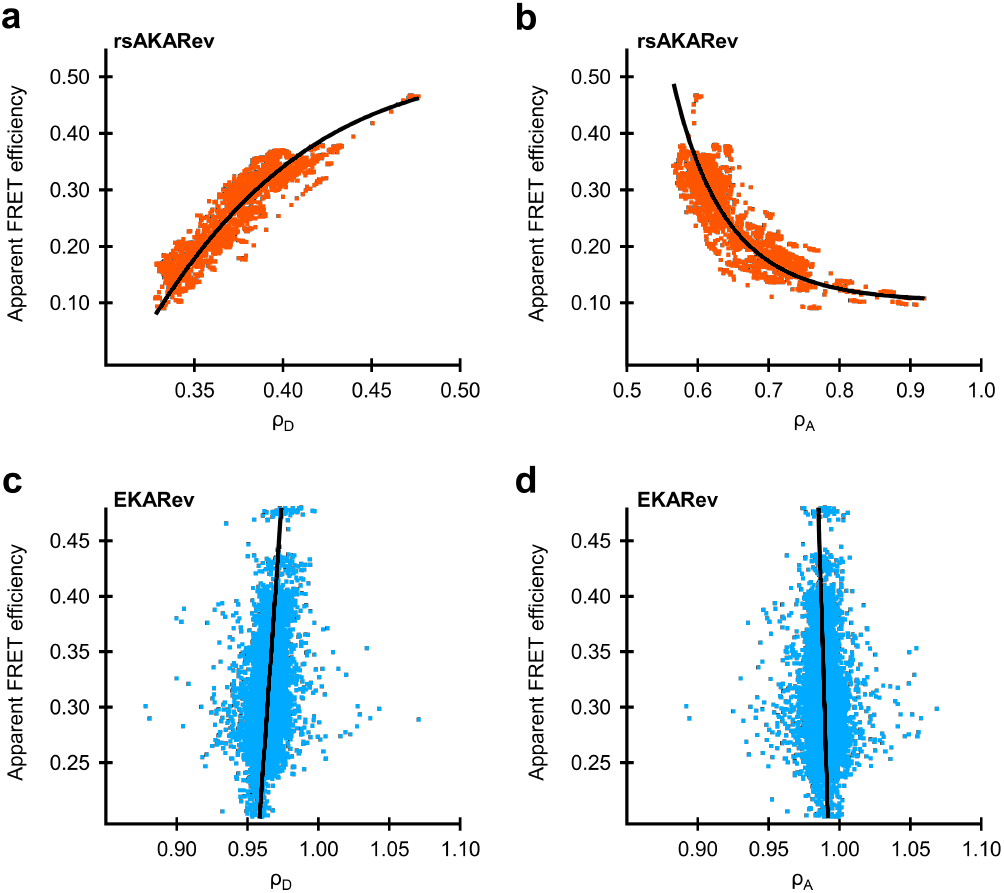
Relation between apparent FRET efficiency and photoswitching ratio of donor emission (a,c) and donor-excited acceptor emission (b,d) as determined by single-sensor experiments on COS-7 cells expressing rsAKARev (orange) or HeLa cells expressing EKARev (blue). *n*=99 (rsAKARev) or 41 (EKARev) cells from 1 experiment. Solid lines represent the empirical fitting functions for rsAKARev (double exponential) or EKARev (straight line) that were used for downstream analysis.

### Simultaneous measurement of two overlapping FRET biosensors

We then focused on the simultaneous measurement of rsAKARev and EKARev expressed within the same cells. Since the emission spectra of both biosensors are essentially indistinguishable, their responses would be difficult to separate based on classical FRET imaging. In order to generate predictable response profiles for rsAKARev and EKARev, we treated the cells sequentially with PMA, Fsk/IBMX, U0126, and H-89. U0126 and H-89 are inhibitors that reverse the responses of the MAPK/ERK and PKA signaling pathways, respectively [17]. Ideally, this sequence of manipulations should result in distinguishable response profiles for rsAKARev and EKARev.

We initially performed these experiments on cells expressing a single biosensor, either rsAKARev, EKARev, or inactive mutants in which the kinase target threonine was mutated to alanine, denoted using a ‘(T/A)’ suffix. We subjected these cells to the measurement scheme shown in Figure 1b, but analyzed the resulting responses in two different ways: either via a conventional intensity-based analysis, using only the ‘on’ fluorescence images acquired after every violet irradiation step (Figure 5a,d,g), or via our full photochromism-based approach (Figure 5b,e,h). Applying both analysis methodologies to the same data makes it possible to detect any discrepancies between the obtained results. Details of the analysis procedure are provided in section 3 of the supplementary information.

**Figure 5:**
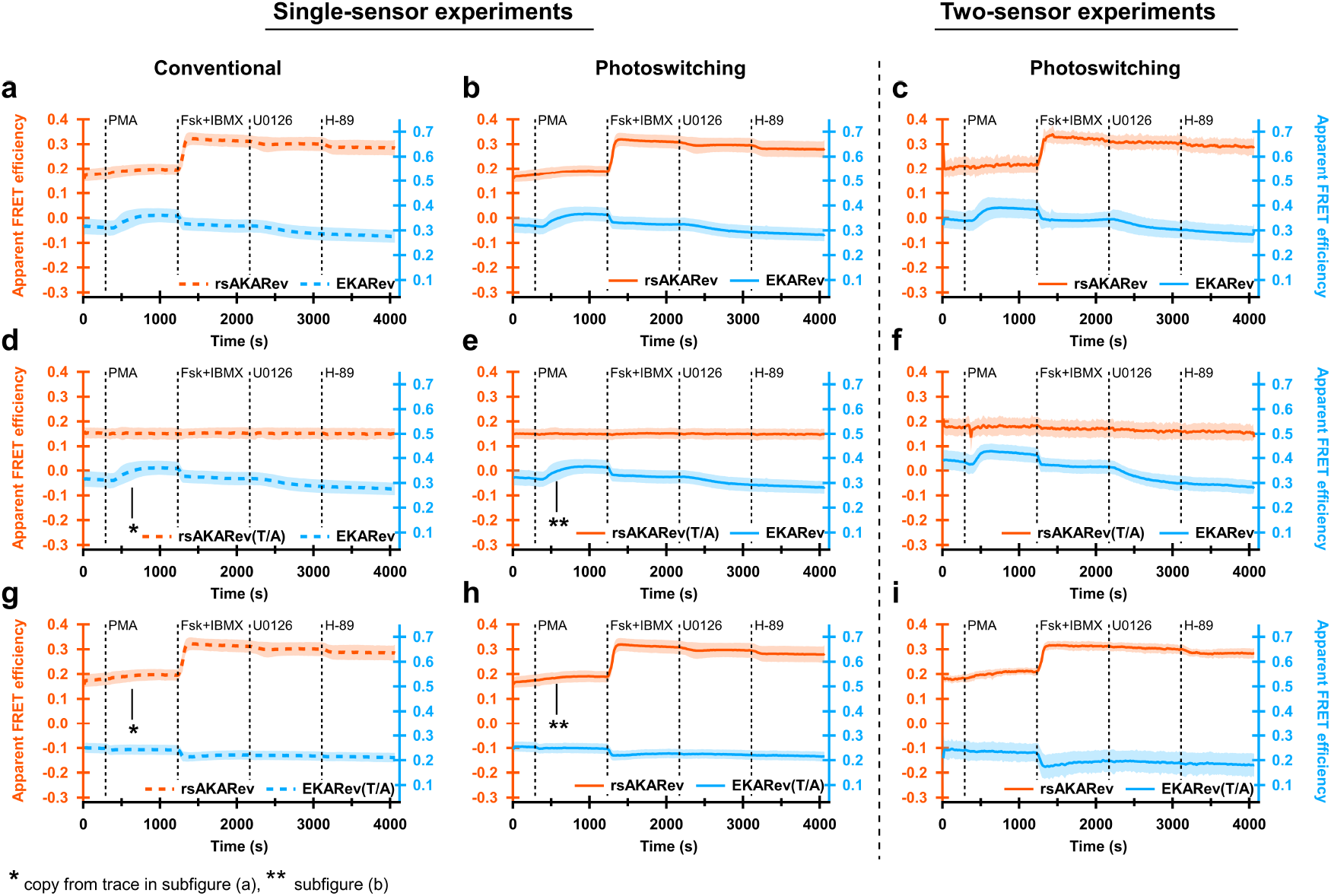
PKA and/or ERK signaling responses in HeLa cells expressing one (left, middle panels) or two (right panels) FRET biosensors. Responses of rsAKARev, rsAKARev(T/A), EKARev and EKARev(T/A) are depicted per column according to the method of analysis. Dashed dark traces indicate results obtained using conventional analysis based on sensitized emission, while solid light traces indicate results obtained using the photochromism-based analysis presented here. All traces represent average apparent FRET efficiencies plus standard deviation (shaded area). *n*=75 (rsAKARev), 132 (EKARev), 12 (rsAKARev(T/A)), 42 (EKARev(T/A)) cells for single-sensor experiments and 80 (rsAKARev+EKARev), 50 (rsAKARev(T/A)+EKARev), 22 (rsAKARev+EKARev(T/A)) cells for two-sensor experiments, from 3, 2, 1, 1 and 2, 1, 1 independent experiments, respectively. Cells were stimulated with PMA (1 μM), forskolin (Fsk, 50 μM) and IBMX (100 μM), U0126 (20 μM) and H-89 (20 μM).

The conventional analysis shows that the biosensors respond as expected to pharmacological treatment (Figure 5a). We did observe an inhibition of the EKARev signal upon addition of Fsk/IBMX, reminiscent of previous reports where it was found that forskolin interferes with EGF-mediated ERK activation [17]. This suggests that a similar mechanism is also at play when stimulating ERK activity with PMA. As expected, the biosensor response of the T/A mutants remained insensitive to the applied sequence of manipulations. Addition of Fsk/IBMX did induce a decrease in the EKARev(T/A) response similar to that observed for EKARev, demonstrating that this response arises from a process other than phosphorylation of the target residue. We also observed a weaker than expected inhibition of the rsAKARev signal upon addition of H-89. However, additional experiments using a fresh sample of H-89 showed the expected reversibility of the biosensor response (Supplementary Figure 2), demonstrating that the limited reversibility found here was due to a poorly performing batch of H-89.

The results from the conventional analysis and our new unmixing strategy are in excellent agreement for the single sensor experiments, reproducing the same features and responses. Taken together, the differences between the photochromism-based and the conventional analysis are small and unlikely to be important for all but the most demanding quantitative experiments.

Having validated our strategy at the level of single biosensors, we then moved on to cells expressing both rsAKARev and EKARev simultaneously. Our analysis reproduces essentially the same trends as observed for the single-sensor case (Figure 5c,f,i), demonstrating that we can indeed separate two FRET pairs that display essentially complete spectral overlap. We do find that the simultaneous imaging of two biosensors leads to a reduction in the signal-to-noise ratio of the separated responses (Supplementary Figures 3 and 4), consistent with the extra processing required to separate the signals, though the cellular responses can still be readily verified.

### Multiplexed detection of three signaling activities

Our separation strategy can be straightforwardly combined with other spectrally-distinct fluorophores or sensors. As a proof-of-concept, we simultaneously expressed rsAKARev and EKARev together with the red-shifted calcium sensor RCaMP [44]. Calcium is known to modulate signaling activities of proteins within both the cAMP/PKA [45, 46] and the MAPK/ERK [47] signaling pathways. Additionally, it has been shown that the PKA-mediated activation of the CREB transcription factor occurs via MAPK/ERK and is mediated by calcium release from intracellular stores [48]. Such inter-pathway dependencies illustrate the potential of multi-parametric biosensing.

We sequentially stimulated the cells with Fsk/IBMX, epidermal growth factor (EGF), ionomycin, and CaCl_2_ in order to provoke distinguishable response profiles. Ionomycin is an ionophore that transports Ca^2+^ ions over lipid bilayers, leading to both calcium release from intracellular stores and calcium uptake from or leakage to the external medium. CaCl_2_ was added as an internal control for the RCaMP response. A schematic overview of the stimulated signaling pathways can be found in Supplementary Figure 5.

Using our strategy, rsAKARev and EKARev responses could be distinguished based on their photochromism, while the red-shifted RCaMP response could be separated based on its different excitation and emission wavelengths. Figure 6a shows the responses of the three biosensors for all 129 analyzed cells. Plots for each individual biosensor can be found in Supplementary Figure 6, together with images of representative cells (Supplementary Figure 7) and their corresponding responses (Supplementary Figure 8). To simplify the following discussion, we also visually indicate three different time intervals during our measurement at the top of Figure 6a.

**Figure 6:**
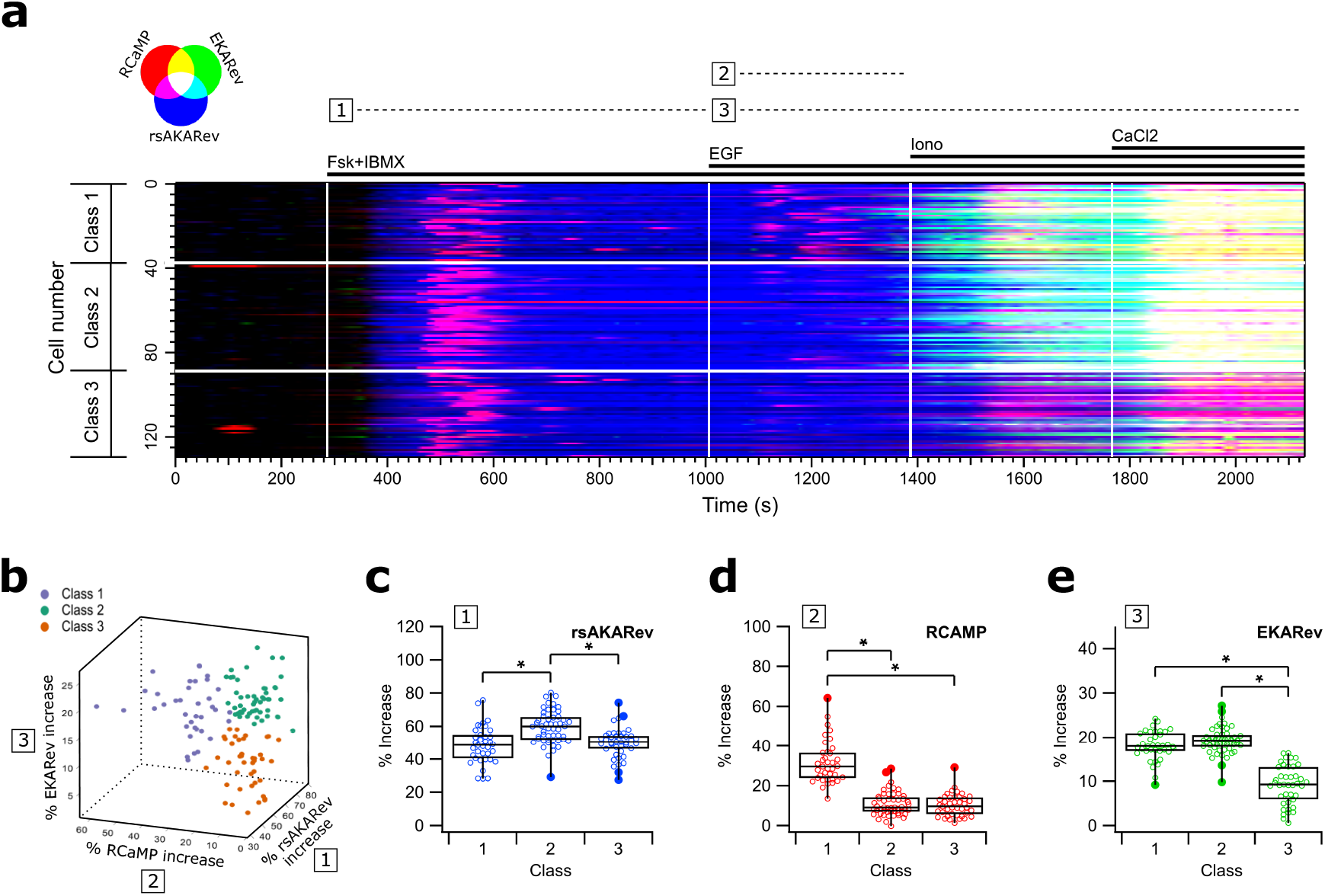
Three biosensor unmixing experiment on HEK293T cells simultaneously expressing rsAKARev, EKARev and RCaMP biosensors. (a) RGB color-coded responses of RCaMP (red), EKARev (green), and rsAKARev (blue). (b) 3D scatter plot of the maximal response, colored by the clustering assignment of each cell. (c,d,e) boxplots of the maximal response observed for rsAKARev (c), RCaMP (d) and EKARev (e) during indicated time-intervals. ‘*’ indicates a significant difference (p<0.01, two-sample t-test following an F-test for equal variances (p<0.05)). *n*=129 cells from 1 experiment. Cells were stimulated with forskolin (Fsk, 50 μM) and IBMX (100 μM), EGF (1 μg/mL), ionomycin (Iono, 10 μM) and CaCl_2_ (2 mM).

As expected, stimulation with Fsk/IBMX (interval 1) strongly increases the PKA activity in all cells, additionally showing that cAMP elevation also mediates a transient calcium response. This is in line with previous work describing cross-talk occurring via PKA-mediated release of calcium from ER-stores [49, 50]. Next, EGF stimulation results in a second release of calcium (interval 2) [51, 52], and in delayed ERK activity (interval 3), with varying responses across cells. The onset of the ERK response overlaps somewhat with the addition of ionomycin, though this response is already initiated before addition in most cells, and has reached a plateau by the time at which the ionomycin-induced release of calcium becomes evident (Supplementary Figures 6 and 8). Taken together, this suggests that ionomycin has a small or no effect on ERK activity.

Visual inspection of the data suggested heterogeneity of the cellular responses (Figure 6a). We quantified the maximal response and timing of the biosensor responses within each interval (see methods, Supplementary Figure 9), and found that k-means clustering could divide the cellular responses into three classes based on their maximal responses, visually shown in Figure 6b and confirmed using partial least squares discriminant analysis (Supplementary Figure 10).

The maximal response of the different sensors showed significant variations between the classes (Figure 6c-e). Fsk/IBMX stimulation appears to have a slightly stronger effect on the PKA activity in cells of class 2, though this does not appear to translate into a higher calcium transient, which was overall very variable. Most striking are the inter-class discrepancies in calcium (interval 2) and ERK (interval 3) signaling upon EGF stimulation. More specifically, cells within classes 1 and 2 respond strongly to EGF-mediated ERK activation, whereas the effect on class 3 cells is rather weak and occurs with an additional delay (Supplementary Figure 9h). Additionally, the sizes of the ERK responses do not appear to be linked to the increase of the preceding calcium release.

These results show the potential of classifying cells based on the heterogeneity in their responses, although it is not immediately clear from the data where this heterogeneity originates from, or if the differences in the signaling activities of the three pathways are biologically correlated in the context of this experiment.

Poor expression of EGF receptors [53, 54] or Fsk/IBMX-induced cAMP-mediated effects [55] could render some cells less sensitive to EGF stimulation, while further heterogeneity may also be contributed by the high levels of variability associated with HEK293 cells [56].

Taken together, our data further supports the notion that the simultaneous imaging of multiple biosensors can readily reveal heterogeneities even in well-established systems, while demonstrating that our scheme can be readily combined with spectrally-orthogonal biosensors to further expand the range of responses that can be measured.

## Conclusions

In this work we set out to separate the contributions of two or more spectrally overlapping FRET pairs based on the photochromism of the donor fluorophores, by monitoring their response to irradiation with off-switching light. We developed a framework to perform the analysis and validated its applicability using numerical simulations. We find that our method can deliver accurate results already at low light levels, though the competition between the FRET process and the photochromism must be characterized for each of the FRET pairs.

We then applied our methodology to FRET-based biosensors, many of which make use of cyan and yellow fluorescent proteins. We showed that the cyan fluorescent protein mTFP0.7 could be used to make a photochromic biosensor for PKA activity, which we named rsAKARev. We combined this sensor with the spectrally overlapping biosensor EKARev, a reporter for ERK kinase activity, and showed that the contributions of both sensors could be readily separated when expressed simultaneously in live cells. Finally, we expanded these experiments to include an additional reporter for Ca^2+^, and showed that this approach could detect cellular heterogeneity in the response to pharmacological stimulation. Our method further illustrates the potential of simultaneously acquiring multiple signaling responses to investigate complex biological networks.

Our method is readily compatible with existing measurement or analysis pipelines, which may make use of the raw emission ratio, sensitized emission ratio, or the calculated FRET efficiency, provided that one characterizes the dependence between this metric and the photoswitching ratio. In practice, this will likely require a separate measurement for each FRET pair where the photoswitching ratio is monitored over varying emission ratios or FRET efficiencies. Such a measurement must only be performed once for each FRET pair and off-switching light dose. The method can be readily integrated with established instrumentation, requiring only the possibility of inserting an off- and an on-switching irradiation in between two FRET acquisitions.

In principle, our procedure can be used to separate the contributions of more than two FRET pairs, provided that the donors are sufficiently separated in terms of their photochromism. Our use of two biosensors also reflects the limited availability of photochromic cyan fluorescent proteins, though many more opportunities are available when considering also photochromic green and red fluorescent proteins, which would be directly compatible with our method.

In summary, we have shown that photochromism of the donor can be leveraged to achieve FRET multiplexing using a simple linear unmixing scheme, on the condition that the responses of the FRET pairs are well characterized. Our strategy can be readily used as part of experiments that require multiplexed interaction sensing and biosensing.

## Acknowledgements

We thank Eva De Jong, Annemie Biesemans, and Fabian Hertel for assistance with the experiments and interpretation, and Hideaki Mizuno (KULeuven) and Alison Tebo (Janelia Research Campus) for insightful suggestions and discussion. We thank Marco Dalla Vecchia for preparing figure 1a. We thank Michiyuki Matsuda (University of Kyoto) for providing a plasmid encoding EKARev. T.R. thanks the Research Foundation Flanders (FWO) for a doctoral fellowship. S.H. thanks the FWO for a postdoctoral fellowship. This work was supported through funding from the Research Foundation Flanders through grants G090819N and G0B8817N, the European Research Council through grant 714688 NanoCellActivity, the KU Leuven through grant C14/17/111, and the National Institutes of Health through grant NIH R01 DK073368 (to J.Z.).

## Author contributions

P.D. designed research. P.D. supervised research, apart from the construction of rsAKARev, which was supervised by J.Z.. T.R., P.D. and F.S. designed experiments. T.R. performed experiments with contributions from F.S.. T.R., W.V. and P.D. developed the analysis methodology. T.R. and S.H. analyzed data. C.S. provided critical input and discussions. T.R. and P.D. wrote the manuscript with input from all authors.

## Methods

### Biosensor construction

The expression plasmids were generated using a combination of polymerase chain reaction (PCR), restriction enzyme digestion and Gibson Assembly molecular cloning techniques. For Gibson Assembly reactions, vector DNA was digested with restriction enzymes cutting the multiple cloning site. All other DNA fragments were generated by PCR with Q5-polymerase using primer sequences with an optimal annealing temperature of 72°C, calculated using the NEB Tm calculator (https://tmcalculator.neb.com), and with a minimal overlap of 20-basepairs between neighboring fragments. All fragments were then purified using gel extraction and subsequently combined in the Gibson Assembly reaction according to manufacturer instructions. rsAKARev was generated by replacing the cDNA for YPet and ECFP in AKAR3EV (pcDNA3.1+ vector) with those for mTFP0.7 and cpVenus172, respectively. For rsAKARev(T/A), PCR primers were modified to introduce a T/A mutation in the PKA substrate region. For the generation of EKARev(T/A), PCR primers were designed to introduce a T/A mutation in the ERK substrate region.

### Cell culture and transfection

HeLa, COS-7 and HEK293T cells were cultured in Dulbecco’s modified Eagle medium (DMEM; Gibco, #31053028) supplemented with 10%(v/v) fetal bovine serum (FBS; Gibco, #70011036), 1X Glutamax (Gibco, #35050061) and 50 μg/ml gentamycin (GM; Sigma-Aldrich, #15750060) and maintained in a humidified incubator (37°C, 5% CO_2_). 24h before transfection, cells were plated onto sterile 35-mm glass-bottom dishes (MatTek) and grown to 30-70% confluence. Cells were subsequently transfected 24-48h before imaging using FuGENE6 transfection reagent (Promega) and plasmid DNA, according to the manufacturer’s protocol. For optimal transfection results, 90 μl DMEM containing 3 μL FuGENE6 was mixed into 10 μL ultrapure water containing a total of 1 μg DNA by pipetting thoroughly and each mixture was left for at least 20 minutes before addition to each dish in dropwise fashion. For multi-biosensor experiments, the relative amounts of plasmid DNA were optimized to obtain comparable expression of all constructs. Cells were then grown for another 24h (HeLa) or 48h (HEK293T, COS-7) before imaging. Cells were finally switched to serum-reduced medium comprising DMEM (HEK293T, COS-7) or FluoroBrite DMEM (Gibco, #A1896702) (HeLa), supplemented with 0.5%(v/v) FBS, 1X Glutamax and 50 μg/ml GM 4-6h (HEK293T) or 16-20h (COS-7, HeLa) before imaging.

### Time-lapse fluorescence imaging

#### Cell treatment

Before imaging, 35-mm glass bottom dishes were washed twice gently with phosphate buffered saline (PBS; Gibco, #70011036) and then imaged in the dark in either calcium-free Hank’s balanced salt solution (HBSS; Gibco, #14185052) supplemented with 0.2%(w/v) bovine serum albumin (BSA; Gibco, #10270106) (HEK293T, COS-7) at room temperature, or in FluoroBrite DMEM supplemented with 0.5% (v/v) FBS and 1X Glutamax (HeLa) at 37°C, 5% CO_2_. Cells were treated at indicated timepoints using a final concentration of 50 μM forskolin (Fsk) (Sigma-Aldrich, #93049), 100 μM 3-isobutyl-1-methylxanthine (IBMX) (Sigma-Aldrich, #410957), 1 μg/mL epidermal growth factor (EGF) (R&D Systems, #236-EG10), 10 μM ionomycin (Iono) (Sigma-Aldrich, #407950), 2 mM CaCl_2_ (Chem-Lab Analytical, #CL05.0371), 1 μM phorbol-12-myristate-13-acetate (PMA) (Sigma-Aldrich, #P1585), 20 μM U0126 (Sigma-Aldrich, #662005) and 20 μM H-89 (Sigma-Aldrich, #B1427), by mixing a 5X concentrated solution, prepared in imaging buffer from freshly thawed DMSO (Fsk, IBMX, PMA, U0126, H-89) or HBSS (CaCl_2_) stocks, into the imaging dish.

#### Equipment

Images were acquired on an Olympus IX-71 inverted microscope coupled to a Spectra X Light Engine (Lumencor), 10× UplanSApo objective (Olympus), ORCA-Flash4.0 LT camera (Hamamatsu), IX2-RFACA motorized fluorescence cube turret (Olympus), Lambda 10-B Optical Filter Changer (Sutter) and H117 high precision motorized stage (Prior). All hardware was controlled using in-house software. Experiments performed at 37°C, 5% CO_2_ were done using a stage-top incubator (H301-NIKON-TI-S-ER, OKOLAB) coupled to a digital CO2 controller unit (CO2-UNIT-BL, OKOLAB).

#### Image acquisition

Following the notation from the theory, the *S*_*DD*_, *S*_*DA*_ and *S*_*AA*_ detection channels were acquired with a T455LP (*S*_*DD*_, *S*_*DA*_) or ZT514RDC (*S*_*AA*_) dichroic mirror and a ET480/40m (*S*_*DD*_) or ET545/40m (*S*_*DA*_, *S*_*AA*_) emission filter (all Chroma). Each sample was excited with blue light (438/29nm, 35.6mW; *S*_*DD*_, *S*_*DA*_) or teal light (513/22nm, 2.45mW; *S*_*AA*_). RCaMP intensity was acquired using a ZT561rdc dichroic and HQ572lp emission filter (all Chroma) with excitation by green light (542/33nm, 39.0mW). All images were acquired during 50 ms camera exposure. Each acquisition was performed during constant time-intervals every 10-30 seconds on multiple positions of the sample.

Irradiation settings for photochromism were determined as the minimum light dose required to reach steady-state fluorescence during ON and OFF switching of mTFP0.7, which was achieved by irradiating 0.2 s with violet light (390/22nm, 26.7mW) and 0.2 s with blue light (438/29nm, 78.35mW), respectively. *S*_*DD*_ and *S*_*DA*_ fluorescence was acquired after each irradiation step; *S*_*AA*_ fluorescence was acquired for only a subset of timepoints to minimize acceptor photobleaching.

#### FRET analysis

Analysis of the fluorescence time-series was performed using in-house analysis software running on Igor Pro 8 (WaveMetrics). For each sample position, multiple regions of interest (ROI) corresponding to single cells were defined, and the average intensity over the shape of the cell was determined at each time-point. Subsequently the intensity was corrected for background by subtracting the average signal of a region outside the cell boundaries. This correction can be improved by subtracting the signal observed for non-transfected cells instead, though we did not pursue this here as the emission from non-transfected cells was much smaller than both the background and the emission from transfected cells. The resulting average intensities were used for downstream analysis.

For the conventional FRET analysis of single-sensor experiments, FRET efficiencies were subsequently calculated according to Equation (6) using only the background-corrected fluorescence detected after each violet irradiation step in the measurement scheme shown in Figure 1b. The photoswitching analysis of single- and two-biosensor experiments was performed as detailed in the theory using Equations (11) to (14). The analysis procedure is described in more detail in supplementary section 3.

From single-sensor experiments, the dependencies *ρ* (Θ) were determined empirically by fitting a double exponential function (rsAKARev) or a straight line (EKARev) on the pooled cell data represented in Figure 4, using orthogonal distance regression.

Coefficients *α*, *δ* and *γ* (Supplementary Table 1) were calculated based on published values for fluorescence quantum yields (QY), instrument parameters, and the available spectral data on the fluorescent protein database FPbase [57], as described in section 4 of the supplementary information.

#### Three-biosensor unmixing experiment

The FRET efficiencies for rsAKARev and EKARev were obtained as described above. The raw RCaMP intensities were obtained by averaging the corresponding emission over the same regions of interest, and also included background removal by subtracting the signal from a region of interest that did not include any cells.

We found that both the FRET and the RCaMP traces showed gradual decreases in amplitude, which we attribute to photobleaching of the fluorophores. We correct for this using a baseline that is determined for every trace.

For FRET measurements, the baseline is determined by fitting the trace with the function

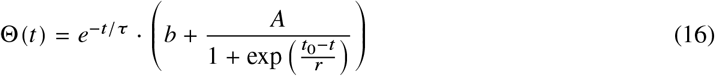

where the factor between parentheses is a standard sigmoidal function. The baseline is then given by *b e*^−*t/τ*^. *τ* is typically much larger than the duration of the experiment.

For RCaMP measurements, we estimated the baseline using by fitting an bi-exponential function to the those parts of the trace that do not show an elevated calcium signal.

We then calculated the responses as

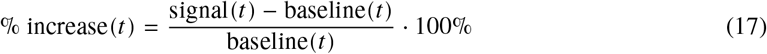

where ‘signal’ is the FRET efficiency for rsAKARev and EKARev, and the fluorescence intensity for RCaMP. The resulting responses are shown in Figure 6a, where we manually optimized the color scaling to highlight the cellular responses. We additionally show example traces, their corresponding baselines and the calculated responses in Supplementary Figures 11 to 13. The cell responses to pharmacological stimulation were quantified by taking the value of the maximum response observed within the time interval following stimulation.

The onset time of the FRET biosensor responses was determined as the time for which a sigmoidal fit of the response increases to 7.6% of its maximum value, while the onset time of the RCaMP data was defined as the timepoint at which the second derivative of the response reached its maximum value.

Partitional k-means clustering was performed in Matlab R2019b based on the responses determined for rsAKARev over time-interval 1, for RCaMP over time-interval 2 and for EKARev over time-interval 3. To validate the classification results of the K-means clustering, partial least squares discriminant analysis (PLS-DA) was performed on the fulltime trace datasets (% increase) assuming the three classes obtained by the K-means clustering. Subsequently, clustering results were cross-validated with the PLS-DA model using a 95% confidence interval. Additional information is available in section 5 of the supplementary information.

#### Statistics and reproducibility

All statistical tests were performed in GraphPad Prism 5 (GraphPad Software). All data was checked for normality using a D’Agostino & Pearson omnibus test. Subsequently, pair-wise comparison was performed using an unpaired two-tailed Student’s t-test or Welch’s unequal variance test for gaussian data, or using the Mann Whitney U test for non-gaussian data with significance set at *P* < 0.01.

## Data Availability

All cell traces and derived datasets can be downloaded from https://doi.org/10.5281/zenodo.4392845.

## Code Availability

All analysis code is available upon reasonable request.

## Notes

### Competing Interest Statement

The authors have declared no competing interest.

