## Supplementary Information for "Simultaneous readout of multiple FRET pairs using photochromism"

### Simultaneous readout of multiple FRET pairs using photochromism
### Supplementary Information

Thijs Roebroek[1], Wim Vandenberg[1], François Sipieter[2], Siewert Hugelier[1], Christophe Stove[3], Jin Zhang[4], Peter Dedecker[1]
[1] Department of Chemistry, KU Leuven, Leuven, Belgium
[2] Université de Paris, Centre National de la Recherche Scientifique, Institut Jacques Monod, Paris, France
[3] Department of Bioanalysis, Ghent University, Ghent, Belgium
[4] Department of Pharmacology, University of California at San Diego, La Jolla, USA


**6 Supplementary figures** **7**

#### 1 Relationship between $F_c$ and $\Theta$

This relationship was detailed already in [1–3], though we here follow a different derivation. It is possible to obtain this from $R_c$ using a relation which we derive as follows. The FRET efficiency ($\Theta$) at a particular timepoint can be expressed using the number of donor excitation cycles that give rise to energy transfer ($n_F$), and the number of donor cycles that do not ($n_D$):

$$\Theta = \frac{n_F}{n_D + n_F} \tag{S.1}$$

$$= \frac{1}{n_D/n_F + 1} \tag{S.2}$$

$n_F$ and $n_D$ can be estimated from the acquired donor ($S_{DD}$) and sensitized acceptor ($F_C$) emission since

$$S_{DD} = n_D \cdot \phi_D \cdot t_D \cdot p_D \cdot \eta \tag{S.3}$$

$$F_C = n_F \cdot \phi_A \cdot t_A \cdot p_A \cdot \eta \tag{S.4}$$

where $\phi_D$ and $\phi_A$ are the fluorescence quantum yields of the donor and acceptor, $t_D$ and $t_A$ are the fractions of the total fluorescence emission of each that are transmitted through the filter and dichroic, $p_D$ and $p_A$ are the probabilities that photons from each will be detected by the camera, and $\eta$ is a parameter that describes the overall collection efficiency of the instrument, which we assume to be identical for both donor and acceptor. Let us now define a correction parameter $\gamma$ [3] given by

$$\gamma = \frac{\phi_A \cdot t_A \cdot p_A}{\phi_D \cdot t_D \cdot p_D} \tag{S.5}$$

which can be estimated using published values for the fluorophore quantum yields as well as specifications from the dichroic, filter, and camera manufacturers. Substitution of Eqs. (S.3) to (S.5) into (S.2) leads to

$$\Theta = \frac{1}{\gamma S_{DD}/F_c + 1} \tag{S.6}$$

providing us with the means to estimate the space and time-resolved FRET efficiency from each $\{S_{DD}, S_{DA}, S_{AA}\}$ dataset.

### 2 Photoswitching ratios for simple FRET pairs

#### 2.1 Dyes that show complete off-switching

Our method requires knowledge of the dependence between the photoswitching ratio $\rho$ and the (sensitized) emission ratio or FRET efficiency. The main text already discussed that for a simple FRET process in which the cross-talk is negligble and only a single species is present, $\rho_D = \rho_A = \rho$. We can further simplify this if we assume that both processes occur independently on account of their difference in time scales, which allows us to treat their relationship using a formalism based on their quantum yields. In particular, we can write:

$$\Theta = \frac{k_{\text{FRET}}}{k_{\text{FRET}} + k_{\text{fl}} + k_{\text{nr}}} \tag{S.7}$$

$$\phi_{sw,0} = \frac{k_{\text{sw}}}{k_{\text{sw}} + k_{\text{fl}} + k_{\text{nr}}} \tag{S.8}$$

$$\phi_{\text{sw}} = \frac{k_{\text{sw}}}{k_{\text{sw}} + k_{\text{fl}} + k_{\text{nr}} + k_{\text{FRET}}} \tag{S.9}$$

where $\phi_{sw,0}$ and $\phi_{\text{sw}}$ are the values of the quantum yield of off-switching in the absence and presence of FRET, respectively, determined by the rate constants for FRET ($k_{\text{FRET}}$), fluorescence ($k_{\text{fl}}$), off-switching ($k_{\text{sw}}$) and all other non-radiative deactivation processes ($k_{\text{nr}}$).

These equations can be reworked to obtain

$$\phi_{\text{sw}} = (1 - \Theta)\, \phi^0_{\text{sw}} \tag{S.10}$$

The photoswitching ratio of the fluorescence is then given by

$$\rho_0 = e^{-k_{\text{exc}} \phi^0_{\text{sw}} T} \tag{S.11}$$

$$\rho = e^{-k_{\text{exc}} \phi_{\text{sw}} T} \tag{S.12}$$

with $T$ the duration of the irradiation and $k_{\text{exc}}$ the excitation rate, which depends on the extinction coefficient and the intensity of the irradiation, but is independent of the occurrence of FRET. Combining Eqs. (S.10) to (S.12) shows that

$$\frac{\ln \rho}{\ln \rho_0} = (1 - \Theta) \tag{S.13}$$

$$\rho = (\rho_0)^{(1-\Theta)} \tag{S.14}$$

which shows that the full dependence is known if one simply measures $\rho_0$ in the absence of FRET, using the same settings for the instrument.

When examined closely, some photochromic molecules do not display complete off-switching but instead reach an equilibrium, caused by competition between the light-induced off-switching and the spontaneous or light-induced on-switching. As we show below, in this case

$$\rho = \frac{(1-\Theta)(\rho_0-\rho_\infty)\left(\frac{\rho_0-\rho_\infty}{1-\rho_\infty}\right)^{\Theta(\rho_\infty-1)}+\rho_\infty}{\Theta\left(\rho_\infty-1\right)+1} \tag{S.15}$$

with $\rho_\infty$ the fractional amount of fluorescence that remains at when the equilibrium is reached, in the absence of FRET. This likewise shows that the full dependence is known by determining $\rho_0$ and $\rho_\infty$ in a control measurement.

#### 2.2 Dyes that show incomplete off-switching

We assume that the photochromism can be described using the simple kinetic scheme

$$\text{Bright} \rightleftharpoons \text{Dark} \tag{S.16}$$

with forward rate constant $k_f$ and reverse rate constant $k_r$. The response of a photochromic molecule to irradiation can then be described as

$$\rho = \frac{k_f e^{-(k_f+k_r)t}+k_r}{k_f+k_r} \tag{S.17}$$

where $\rho$ is the photoswitching ratio as described in the main text. The equilibrium is reached for $t\to\infty$ and corresponds to

$$\rho_\infty = \frac{k_r}{k_f+k_r} \tag{S.18}$$

In the absence of FRET, this equilibrium corresponds to

$$\rho_\infty = \frac{k_r}{\alpha\phi_0+k_r} \tag{S.19}$$

where we here use $\alpha$ to denote the excitation rate and $\phi_0$ is the quantum yield of switching. Furthermore $k_f = \alpha\phi$ in the presence of FRET. Solving equation (S.19) for $k_r$ and substituting this into equation (S.17) leads to

$$\rho_0 = \frac{\alpha\phi_0 e^{t\left(\frac{\alpha\rho_\infty\phi_0}{\rho_\infty-1}-\alpha\phi_0\right)}-\frac{\alpha\rho_\infty\phi_0}{\rho_\infty-1}}{\alpha\phi_0-\frac{\alpha\rho_\infty\phi_0}{\rho_\infty-1}} \tag{S.20}$$

$$\rho = \frac{\alpha\phi e^{t\left(\frac{\alpha\rho_\infty\phi_0}{\rho_\infty-1}-\alpha\phi\right)}-\frac{\alpha\rho_\infty\phi_0}{\rho_\infty-1}}{\alpha\phi-\frac{\alpha\rho_\infty\phi_0}{\rho_\infty-1}} \tag{S.21}$$

in the absence and presence of FRET, respectively. Both equations can be reworked to remove the exponential, leading to

$$\log\left(\frac{\rho_0\left(\alpha\phi_0-\frac{\alpha\rho_\infty\phi_0}{\rho_\infty-1}\right)+\frac{\alpha\rho_\infty\phi_0}{\rho_\infty-1}}{\alpha\phi_0}\right) = t\left(\frac{\alpha\rho_\infty\phi_0}{\rho_\infty-1}-\alpha\phi_0\right) \tag{S.22}$$

$$\log\left(\frac{\rho\left(\alpha\phi-\frac{\alpha\rho_\infty\phi_0}{\rho_\infty-1}\right)+\frac{\alpha\rho_\infty\phi_0}{\rho_\infty-1}}{\alpha\phi}\right) = t\left(\frac{\alpha\rho_\infty\phi_0}{\rho_\infty-1}-\alpha\phi\right) \tag{S.23}$$

Dividing both equations then yields

$$\frac{\phi-\rho_\infty\phi}{\phi_0}+\rho_\infty = \frac{\log\left(\rho-\frac{(\rho-1)\rho_\infty\phi_0}{(\rho_\infty-1)\phi}\right)}{\log\left(\frac{\rho_\infty-\rho_0}{\rho_\infty-1}\right)} \tag{S.24}$$

Substituting $\phi = \phi_0(1-\Theta)$:

$$\frac{(1-\Theta)\phi_0 - (1-\Theta)\rho_\infty\phi_0}{\phi_0} + \rho_\infty = \frac{\log\left(\rho - \frac{(\rho-1)\rho_\infty}{(1-\Theta)(\rho_\infty-1)}\right)}{\log\left(\frac{\rho_\infty-\rho_0}{\rho_\infty-1}\right)} \tag{S.25}$$

which can be reworked to obtain the final result

$$\rho = \frac{(\Theta-1)(\rho_\infty-\rho_0)\left(\frac{\rho_\infty-\rho_0}{\rho_\infty-1}\right)^{\Theta(\rho_\infty-1)} + \rho_\infty}{\Theta(\rho_\infty-1)+1} \tag{S.26}$$

### 3 Details of the analysis procedure

Each experiment consists of a series of fluorescence images acquired in a time lapse. On each iteration of the time lapse (timepoint) we record the fluorescence in the $S_{DD}$, $S_{DA}$, and $S_{AA}$ channels. Furthermore, $S_{DD}$ and $S_{DA}$ are each measured after irradiation with on-switching light ($S_{DD,\text{on}}$, $S_{DA,\text{on}}$) and after irradation with off-switching light ($S_{DD,\text{off}}$, $S_{DA,\text{off}}$), such that the photochromism of the system can be probed (illustrated in figure 1 in the main text). There is no modulation in the $S_{AA}$ emission, which is therefore only acquired after on-switching irradiation. In practice we only acquire the $S_{AA}$ channel for a subset of the measurement points since this signal varies only slowly.

We then extract a number of intensity trajectories from these images:

1. The acquired data consists of images showing cells and background regions. We first define a region-of-interest (ROI) for each cell in the field-of-view. These regions can be reused for the entire experiment or can be updated during the measurement to adapt to motion of the cells. One could also define separate ROIs for smaller features of the cell, or apply our analysis on a per-pixel basis.

2. The pixels in each ROI are averaged together for each image in the stack. The result is the raw $S_{DD}$, $S_{DA}$, and $S_{AA}$ intensities at each timepoint for each channel and for each cell in the experiment.

3. The raw $S_{DD}$, $S_{DA}$, and $S_{AA}$ intensity time traces are then corrected for background emission by subtracting the signal from a background ROI in the corresponding images at each respective timepoint. This background ROI can consist of a cell-free region, or one or more non-transfected cells if the signal from such cells is not negligible.

The resulting data can then be analyzed purely based on the acquired intensities, which we have described as the 'conventional analysis' in the manuscript, or using the photochromism of the FRET donor. We detail both of these approaches in what follows.

#### 3.1 Conventional FRET analysis

For this analysis we only use the signals recorded immediately after on-switching of the donor. The result of this analysis can be expressed in a number of ways, depending on how rigorously the data is corrected:

- using the raw emission ratio $R$. This is calculated using $S_{DD,\text{on}}$ and $S_{DA,\text{on}}$ following Eq. (3).
- using the sensitized emission ratio $R_c$:
  1. calculate $F_c$ using $S_{DD,\text{on}}$, $S_{DA,\text{on}}$, $S_{AA}$ and correction factors $\alpha$ and $\delta$ following main text Eq. (4).
  2. calculate $R_c$ using $F_c$ and $S_{DD,\text{on}}$ following main text Eq. (5).
- using the FRET efficiency $\Theta$ by first calculating $F_c$ and then introducing this into main text Eq. (6).

The conventional analysis cannot take advantage of the photochromism of the donor fluorophore. If off-switching light is applied then this analysis is only performed using the 'on' fluorescence images. The 'off' fluorescence images are not used.

#### 3.2 Photochromism-based FRET analysis

The photochromism-based analysis can be performed whenever one of the FRET donors shows photochromic behavior. We assume that the sample is labeled with two different FRET pairs that differ in their photochromism, though the same procedure can be generalized to the separation of three or more FRET pairs.

The input data for each timepoint in an intensity trace consists of $S_{DD,\text{on}}$, $S_{DD,\text{off}}$, $S_{DA,\text{on}}$, $S_{DA,\text{off}}$, and $S_{AA}$, where 'on' and 'off' refer to the intensities measured before and after light induced off-switching (see main text Fig. 1). Our goal is to identify the contributions and FRET efficiencies of both pairs that best fit this measured data.

We also require knowledge of the relationship between the photochromism and the efficiency of the energy transfer for each of the FRET pairs, meaning that we need to know the values of main text equations (7) and (8) for all observed FRET efficiencies. In practice, this comes down to either applying a theoretical model such as main text Eq. (10) or performing empirical measurements on systems expressing just a single FRET pair (e.g. main text Fig. 4). In these measurements the system is stimulated such that the FRET efficiencies varies over a range of values, and $\rho_D$ and $\rho_A$ are determined for each of the observed FRET efficiencies. The resulting set of $\{\Theta, \rho_D\}$ and $\{\Theta, \rho_A\}$ measurements can then be visualized in a scatter plot and fitted to an appropriate model that captures the observed variation. This fitting should be performed with orthogonal distance regression or a similar method that recognizes that there are measurement errors not only in the $y$ but also in the $x$ variables. The separation procedure itself can deliver the uncorrected emission ratio $R$ for each FRET pair, or the corrected emission ratio $R_c$ or $\Theta$, as required by the user.

##### 3.2.1 Separating based on $R$

In this case our goal is to find the uncorrected emission ratios $R_1$ and $R_2$ associated with both FRET pairs. The experimental data consists of $S_{DD,\text{on}}$, $S_{DD,\text{off}}$, $S_{DA,\text{on}}$, and $S_{DA,\text{off}}$ for every timepoint and ROI. There is no need to acquire the $S_{AA}$ channel since no corrections are performed, though this does mean that the analysis is less robust against variations in donor-acceptor stoichiometry.

We analyze the data using nonlinear fitting. At every timepoint and ROI the experimental data are fitted using a function that calculates the following values:

$$S_{DD,\text{on}} = S_{DD,\text{on},1} + S_{DD,\text{on},2} \tag{S.27}$$

$$S_{DD,\text{off}} = \rho_{D,1}\, S_{DD,\text{on},1} + \rho_{D,2}\, S_{DD,\text{on},2} \tag{S.28}$$

$$S_{DA,\text{on}} = R_1\, S_{DD,\text{on},1} + R_2\, S_{DD,\text{on},2} \tag{S.29}$$

$$S_{DA,\text{off}} = \rho_{A,1}\, R_1\, S_{DD,\text{on},1} + \rho_{A,2}\, R_2\, S_{DD,\text{on},2} \tag{S.30}$$

where the fit optimizes the values of $S_{DD,\text{on},1}$, $S_{DD,\text{on},2}$, $R_1$, and $R_2$. The values for the switching ratios are recalculated for each iteration of the fit based on the current guesses for $R_1$ and $R_2$ and the knowledge of the relation between the photochromism and energy transfer. The fit terminates when the calculated $S_{DD,\text{on}}$, $S_{DD,\text{off}}$, $S_{DA,\text{on}}$, and $S_{DA,\text{off}}$ best match the measured values. In practice we run this fit in Igor Pro using the 'FuncFit' operation, though this can be implemented in any software that allows the nonlinear fitting of custom fit functions.

The fit terminates when $S_{DD,\text{on}}$, $S_{DD,\text{off}}$, $S_{DA,\text{on}}$, and $S_{DA,\text{off}}$ best approximate the measured values.

##### 3.2.2 Separating based on $R_c$ or $\Theta$

In this case our goal is to find the absolute FRET efficiencies or $R_c$ values associated with the two FRET pairs. The experimental data constists of the $S_{DD,\text{on}}$, $S_{DD,\text{off}}$, $S_{DA,\text{on}}$, $S_{DA,\text{off}}$, and $S_{AA}$ values at every timepoint and for every ROI. The fit function calculates the following values:

$$S_{DD,\text{on}} = S_{DD,\text{on},1} + S_{DD,\text{on},2} \tag{S.31}$$

$$S_{DD,\text{off}} = \rho_{D,1}\, S_{DD,\text{on},1} + \rho_{D,2}\, S_{DD,\text{on},2} \tag{S.32}$$

$$F_{C,\text{on},1} = \gamma_1 \frac{S_{DD,\text{on},1}}{(1/\Theta_1 - 1)} \tag{S.33}$$

$$F_{C,\text{on},2} = \gamma_2 \frac{S_{DD,\text{on},2}}{(1/\Theta_2 - 1)} \tag{S.34}$$

$$S_{DA,\text{on}} = F_{C,\text{on},1} + F_{C,\text{on},2} + \alpha_1\, S_{DD,\text{on},1} + \alpha_2\, S_{DD,\text{off},2} + \beta S_{AA} \tag{S.35}$$

$$S_{DA,\text{off}} = \rho_{A,1}\, F_{C,\text{on},1} + \rho_{A,2}\, F_{C,\text{on},2} + \alpha_1\, \rho_{D,1}\, S_{DD,\text{on},1} + \alpha_2\, \rho_{D,2}\, S_{DD,\text{on},2} + \beta\, S_{AA} \tag{S.36}$$

where the fit optimizes the values of $S_{DD,\text{on},1}$, $S_{DD,\text{on},2}$, $\Theta_1$, $\Theta_2$. We can also elect to leave $S_{AA}$ as parameter to be optimized by the fit if its value is affected by noise. We also note that the values for the switching ratios are recalculated for each iteration of the fit based on the current guesses of $\Theta_1$ and $\Theta_2$ and the knowledge of the relation between the photochromism and energy transfer.

The fit terminates when the calculated $S_{DD,\text{on}}$, $S_{DD,\text{off}}$, $S_{DA,\text{on}}$, $S_{DA,\text{off}}$, and $S_{AA}$best match the measured values. The convergence of the fit can be improved by imposing physical constraints on the optimization procedure, such as that $0 \leq \Theta \leq 1$. In practice we run this fit in Igor Pro using the 'FuncFit' operation, though this can be implemented in any software that allows the nonlinear fitting of custom fit functions.

Regardless of the approach taken, the fitting is repeated for every timepoint in the measurement, leading to the time trajectories shown in, for example, main text Fig. 5.

### 4 Determination of correction factors $\alpha$, $\delta$ and $\gamma$

Supplementary Table 1 summarizes the correction factors that were used to calculate the apparent FRET efficiencies using Eqs. (4) and (6). In these, $\alpha$, $\delta$ and $\gamma$ were determined as follows:

$$\alpha = \frac{Em_D^{DA}}{Em_D^{DD}} \tag{S.37}$$

$$\delta = \frac{I_D Ex_A^{DA}}{I_A Ex_A^{AA}} \tag{S.38}$$

$$\gamma = \frac{\phi_A Em_A^{DA}}{\phi_D Em_D^{DD}} \tag{S.39}$$

where $Em$ is the collection efficiency of the emission light and $Ex$ the excitation efficiency of the donor (D) or acceptor (A), in the respective detection channel ($S_{DD}$, $S_{DA}$, $S_{AA}$). $\phi$ is the reported fluorescence quantum yield for donor (D) and acceptor (A) and $I$ is the intensity of the light at the sample when using donor (D) or acceptor (A) excitation wavelengths. Quantum yields were used as reported for mTFP0.7 [4], ECFP [5] and YPet[6] and for cpVenus172 approximated by that of YPet. Spectral data and instrument parameters were obtained from the database FPBase [7]. Spectral data for mTFP0.7 and cpVenus172 were approximated by those of mTFP1 [4] and YPet [6], respectively.

Table 1: Correction parameters used to determine FRET efficiencies. The same parameters were used for analyzing the phosphorylation insensitive (T/A) mutants.

| | rsAKARev | EKARev |
|---|---|---|
| $\alpha$ | 0.40 | 0.42 |
| $\delta$ | 0.1 | 0.1 |
| $\gamma$ | 2.0 | 2.6 |

### 5 K-means clustering and PLS-DA

#### 5.1 K-means clustering

In order to link the observed signaling activities for PKA, ERK and calcium, we first determined the maximal response (Supplementary Fig. 9a,c,g) (see methods) over three time-intervals (as indicated in Fig. 6a) that characterize the responses of each biosensor to cell treatment with Fsk/IBMX or EGF. These maximal responses were taken together in a reduced dataset (three values for each cell), and were subsequently used for data clustering according to a K-means algorithm [8].

This algorithm is an unsupervised method that partitions this dataset into $K$ distinct (and non-overlapping) clusters by iteratively (re)assigning the different data points to a single cluster and minimizing the distance between each datapoint and the cluster centroid. Here, the number of clusters is pre-defined, and for this specific dataset K-means clustering based on two, three and four classes was investigated, with an optimal clustering resulting from using three classes.

#### 5.2 PLS-DA

To confirm the three classes/clusters obtained from the reduced dataset, a partial least squares - discriminant analysis (PLS-DA) model [9, 10] was calculated using the full time-trace datasets (responses shown in Fig. 6a). This PLS-DA model was calculated by maximizing the inter-class variances for three classes, by rotating the dataspace along three latent variables (LVs), which comprise a linear combination of variables from the original data dataspace (the full timetrace responses).

A PLS-DA model with three LVs, determined by a leave-one-out cross validation and effectively reducing the size of the data set with a factor 10 000, is used to validate the class assigned by the K-means clustering method to each data point. The model (Supplementary Fig. 10a-c) classifies most data points reasonably well with an acceptable number of misclassifications (~10%).

Projecting the datapoints obtained from the K-means clustering on the 3-LV PLS-DA model shows that all three classes effectively belong to a different subspace, with minimal overlap between these subspaces. Class 3 (blue upward triangles in Supplementary Fig. 10d) can be mainly separated from class 1 and class 2 by LV3, while the separation between class 1 (green squares in Fig. 10d) and class 2 (cyan downward triangles in Supplementary Fig. 10d) is mainly established by LV 1 and LV 2. No further interpretation of these LVs was performed as PLS-DA was only used as a tool to validate the obtained classes by the K-means clustering algorithm on the reduced data set.

### 6 Supplementary figures

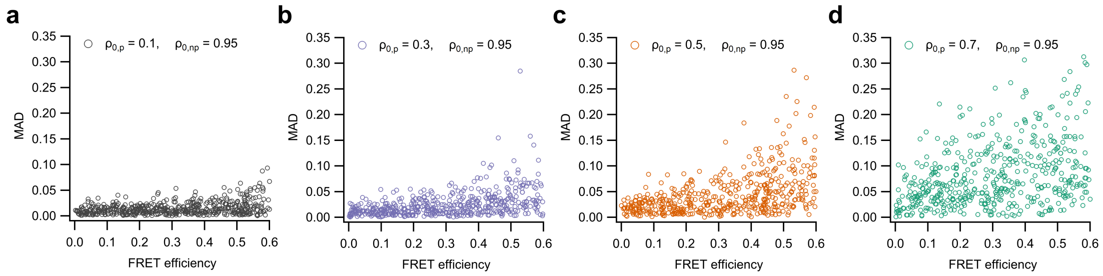

Supplementary Figure 1: Scatter plots showing the analysis performance (mean absolute deviation (MAD)) on simulated data for different FRET efficiencies and photoswitching ratios of the photochromic ($\rho_{0,p}$) and non-photochromic ($\rho_{0,np}$) donor. The conditions for $\rho_{0,p}$ and $\rho_{0,np}$ are shown in the legend of each subfigure.

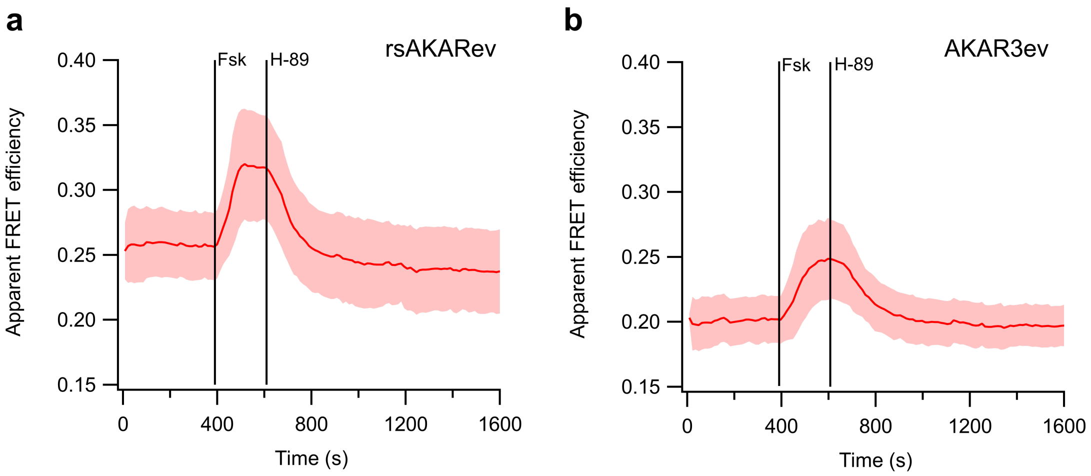

Supplementary Figure 2: PKA activity measured in HeLa cells expressing biosensors rsAKARev (a) or AKAR3ev (b) during stimulation and inhibition. Full lines indicate the average apparent FRET efficiency; shaded area is the standard deviation. Cells were stimulated with forskolin (Fsk, 50 µM) followed by inhibition with H-89 (20 µM). n=19 (rsAKARev) or 48 (AKAR3ev) cells from two independent experiments each. Data was acquired following the irradiation scheme in Fig. 1b and analyzed by conventional analysis. AKAR3ev was analyzed using the same correction factors as those for EKARev.

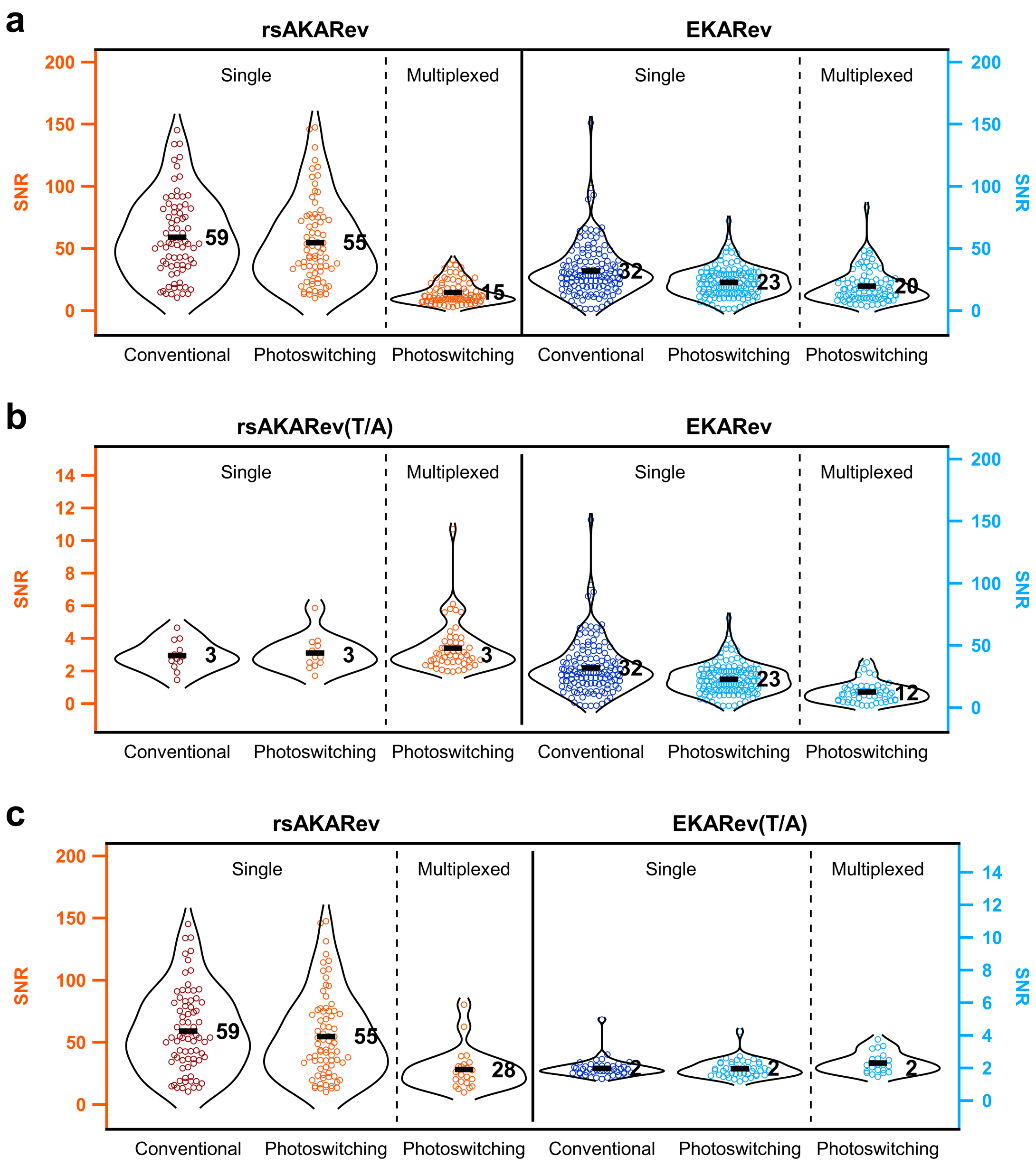

Supplementary Figure 3: Violinplots showing the signal-to-noise (SNR) of all individual responses used to create Fig. 5 of the main text. (a), (b) and (c) correspond to the data shown in Fig. 5a,b,c, Fig. 5d,e,f and Fig. 5g,h,i, respectively. rsAKARev and rsAKARev(T/A) are shown on left axis; EKARev and EKARev(T/A) on the right axis. Black bar and number indicate the mean SNR. SNR was calculated from each individual FRET response time-course by dividing the maximum apparent FRET efficiency by the standard deviation of the baseline before stimulation.The SNR for the T/A mutants is near-zero since these variants are not affected by the pharmacological treatment.

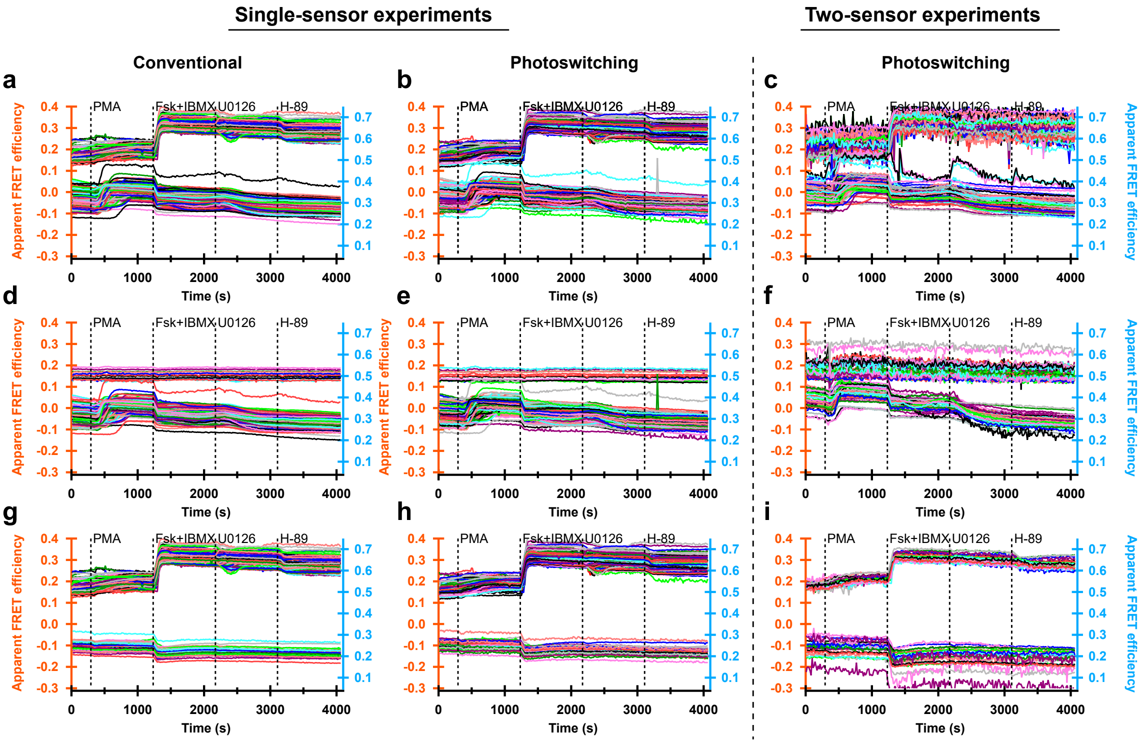

Supplementary Figure 4: Individual apparent FRET efficiency time-traces of the cells that were included in main text Fig. 5. Within each subfigure, each individual trace was assigned a random color to enhance contrast between the different traces. *n*=75 (rsAKARev), 132 (EKARev), 12 (rsAKARev(T/A)), 42 (EKARev(T/A)) cells for single-sensor experiments and 80 (rsAKARev+EKARev), 50 (rsAKARev(T/A)+EKARev), 22 (rsAKARev+EKARev(T/A)) cells for two-sensor experiments, from 3, 2, 1, 1 and 2, 1, 1 independent experiments, respectively. Cells were stimulated with PMA (1 μM), forskolin (Fsk, 50 μM) and IBMX (100 μM), U0126 (20 μM) and H-89 (20 μM).

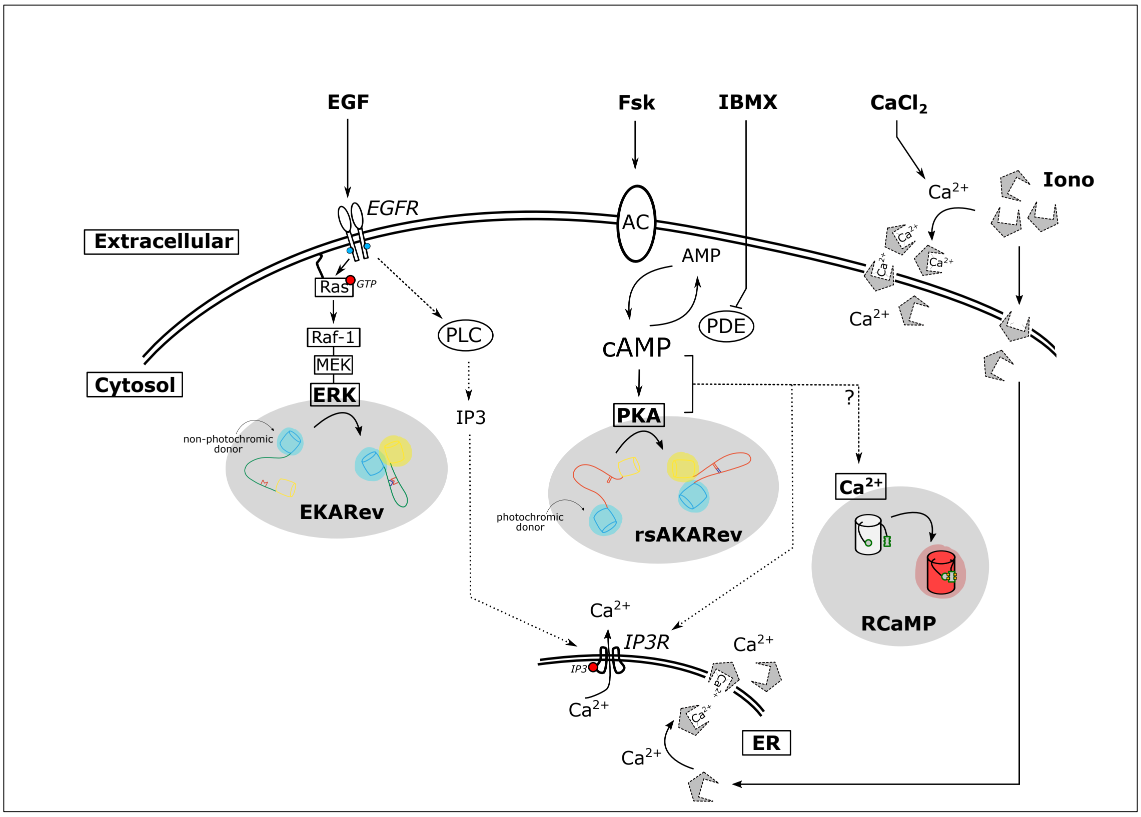

Supplementary Figure 5: Schematic overview of the biochemical signaling activities that are observed during stimulation in Fig. 6. Canonical signaling pathways activated by provided pharmacological compounds are indicated by full lines. Plausible pathways associated with the crosstalk perceived during the experiment are indicated by dashed lines. Markers at the end of each line indicate whether the effect is stimulating or inhibiting. In summary: forskolin (Fsk) and 3-isobutyl-1-methylxanthine (IBMX) elevate cAMP levels via activation of adenylyl cyclase (AC) and inhibition of phosphodiesterases (PDE), respectively, in turn resulting in increased PKA activity. Additionally, cAMP and/or PKA activation mediates calcium release, possibly via the sensitization of Ins3P receptors (IP3R) to basal Ins3P levels [11, 12] or otherwise via unidentified mechanisms [13, 14]. Next, epidermal growth factor (EGF) stimulates its receptor (EGFR) and the downstream MAPK/ERK signaling cascade. EGFR activity additionally results in phospholipase C$\gamma$ (PLC) activation and Ins3P production [15, 16], resulting in calcium release from internal stores via IP3R. Ionomycin (Iono) is an ionophore that transports $Ca^{2+}$ ions over lipid bilayers, leading to both calcium release from intracellular stores (e.g. ER) and calcium uptake from or leakage to the external medium. $CaCl_2$ addition further increases the available calcium for uptake from the external medium.

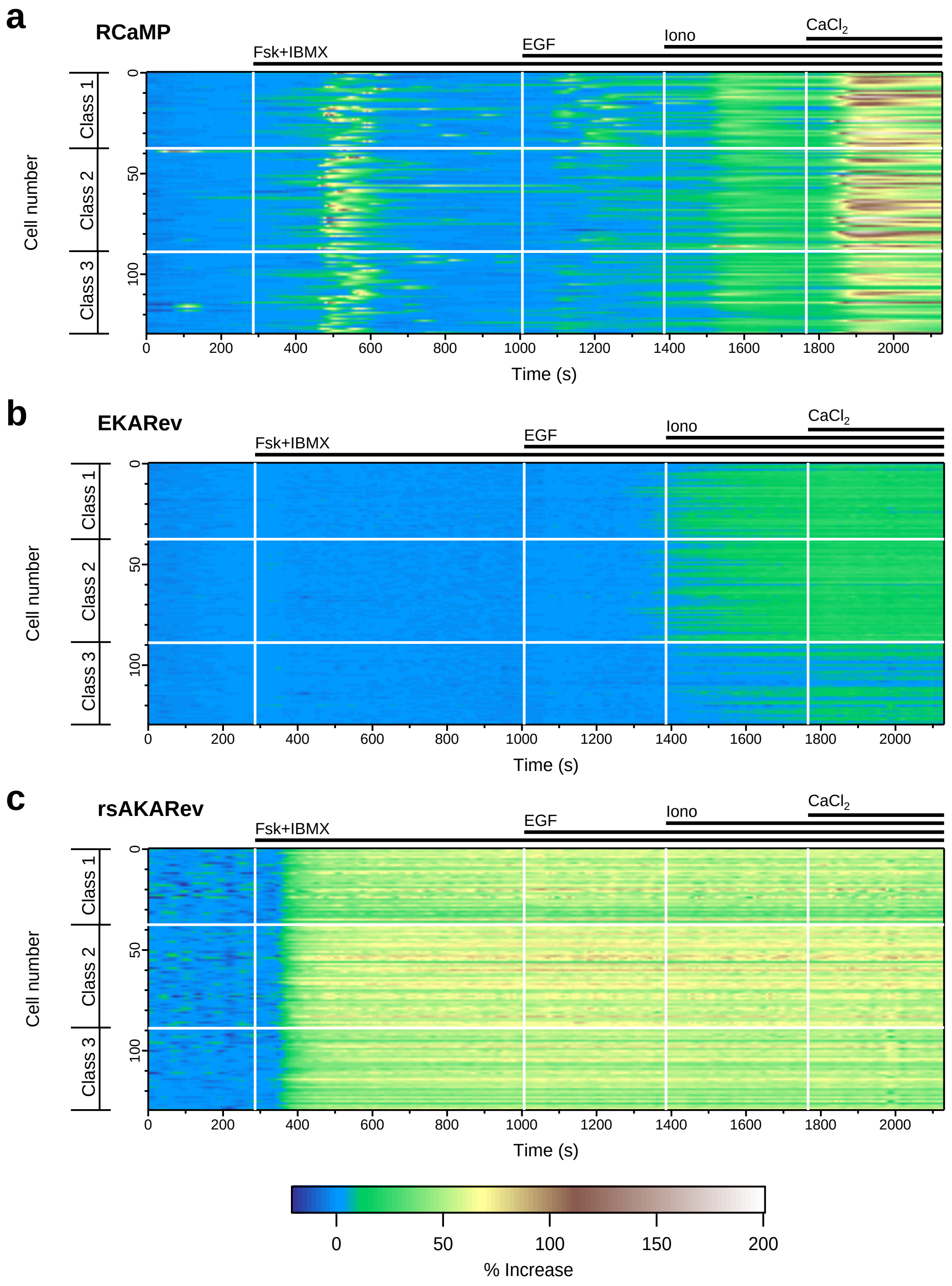

Supplementary Figure 6: 2D plots showing the responses (% increase) of RCaMP (a), EKARev (b) and rsAKARev (c) for all 129 analyzed HEK293T cells expressing the three biosensors simultaneously. Colors were scaled from the minimum to maximum over all measured responses for the three biosensors.

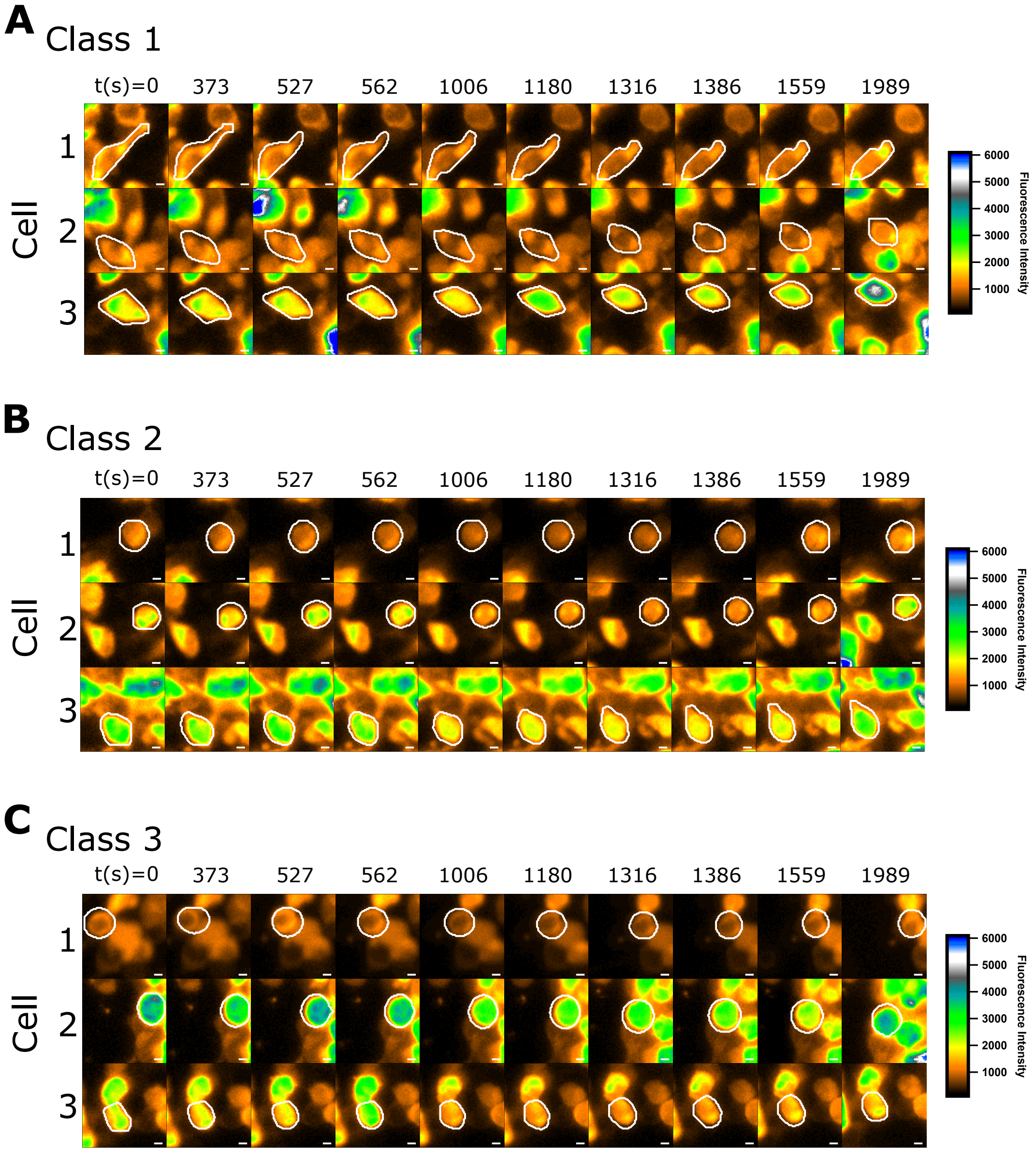

Supplementary Figure 7: Example images of representative HEK293T cells expressing RCaMP, rsAKARev and EKARev. (a-c) Show the raw RCaMP fluorescence intensity images of cells from class 1-3, respectively at the indicated time-points. Time-points represent relevant changes in the responses of RCaMP, rsAKARev and EKARev. The boundaries of the regions of interest at each specific timepoint that were used to designate the cell area are shown in white. Scale bar, 5 µm.

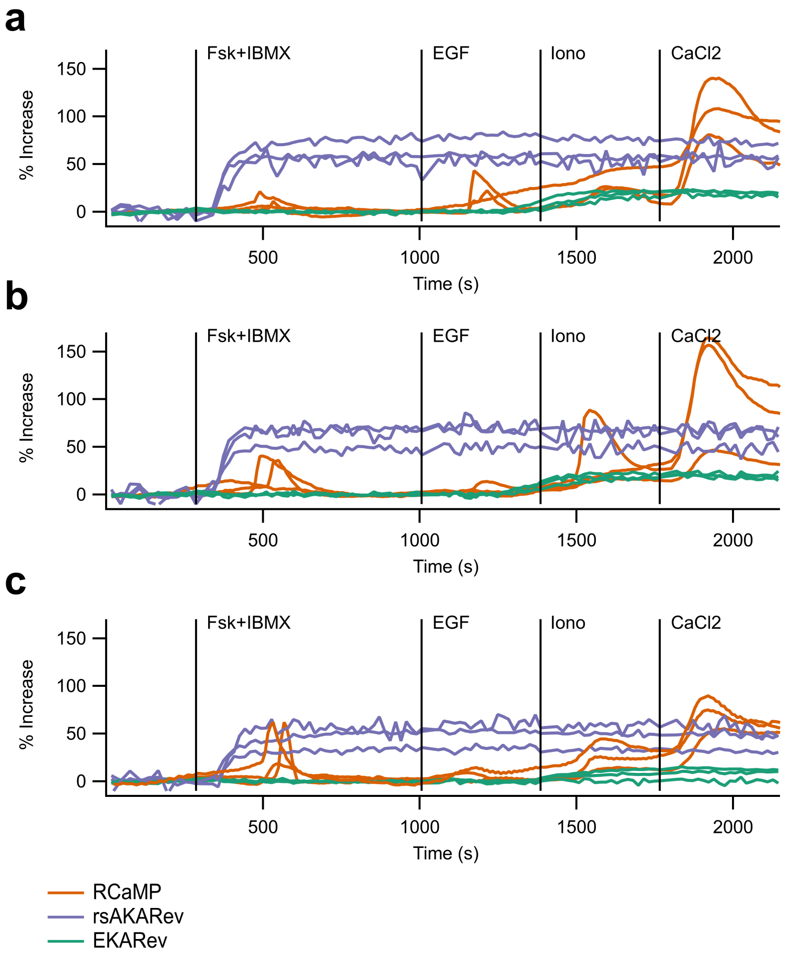

Supplementary Figure 8: Example responses of respresentative HEK293T cells simultaneously expressing RCaMP, rsAKARev and EKARev. (a-c) show the % increase (see Methods) of RCaMP (left axis), rsAKARev (right axis) and EKARev (right axis) corresponding to cells assigned to class 1-3, respectively (cfr. Supplementary Fig. 7). Timepoints of pharmacological stimulation are indicated by vertical lines. Cells were stimulated with forskolin (Fsk, 50 µM) and IBMX (100 µM), EGF (1 µg/mL), ionomycin (Iono, 10 µM) and $CaCl_2$ (2 mM).

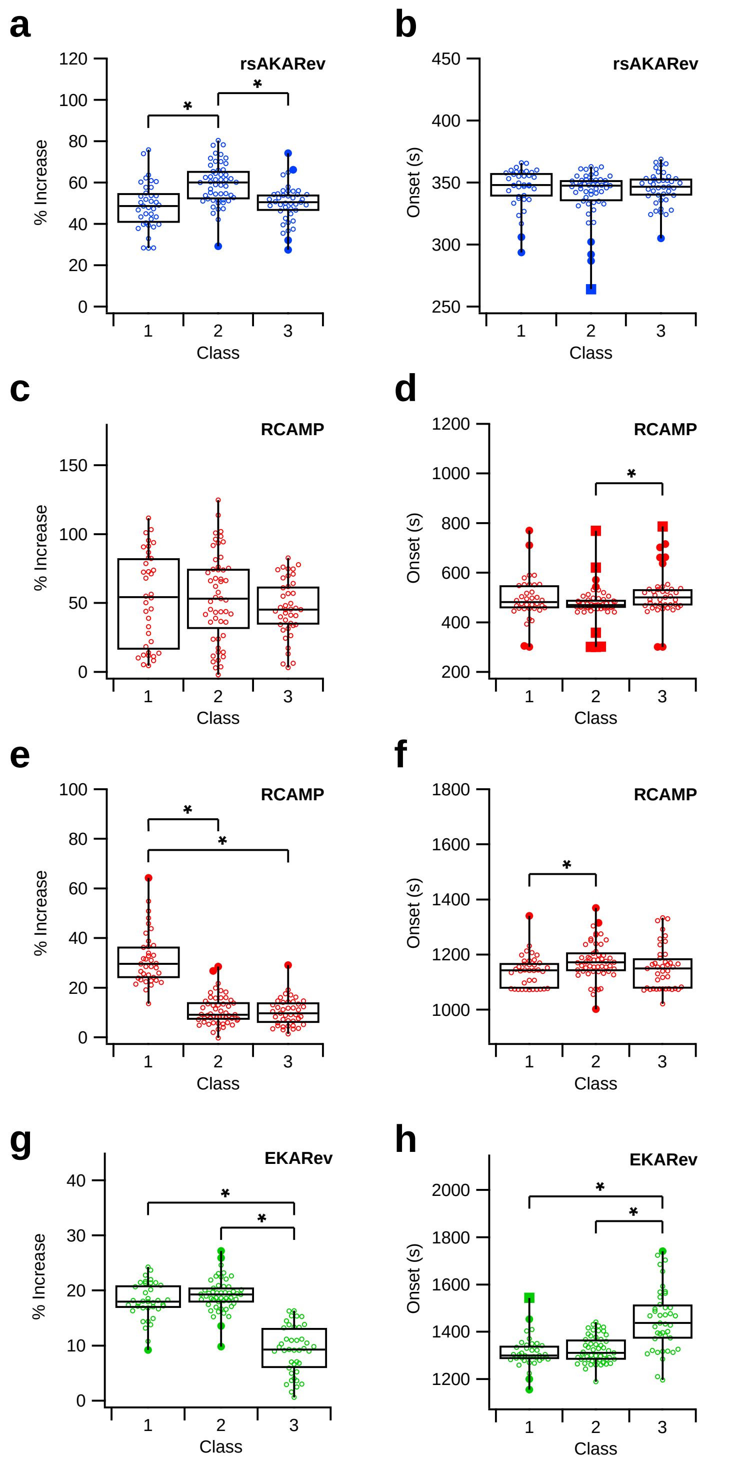

Supplementary Figure 9: Boxplots showing increase and timing of responses in HEK293T cells expressing RCaMP, rsAKARev and EKARev. Each point represents the maximal response or onset of one cell, categorized per class. (a-b, blue) and (c-d, red) show rsAKARev increase (a), rsAKARev onset (b), RCaMP increase (c) and RCaMP onset (d) after addition of Forskolin (Fsk; 50 µM) and IBMX (100 µM) (interval 1 in Fig. 6a). (e-f, red) show RCaMP increase (e), RCaMP onset (f) after addition of EGF (1 µg/ml) (interval 2 in Fig. 6a). (g-h, green) show EKARev increase (g) and EKARev onset (h) after addition of EGF (1 µg/ml) (interval 3 in Fig. 6a). * indicates a significant difference ($p<0.01$, two-sample t-test following an F-test for equal variances ($p<0.05$))

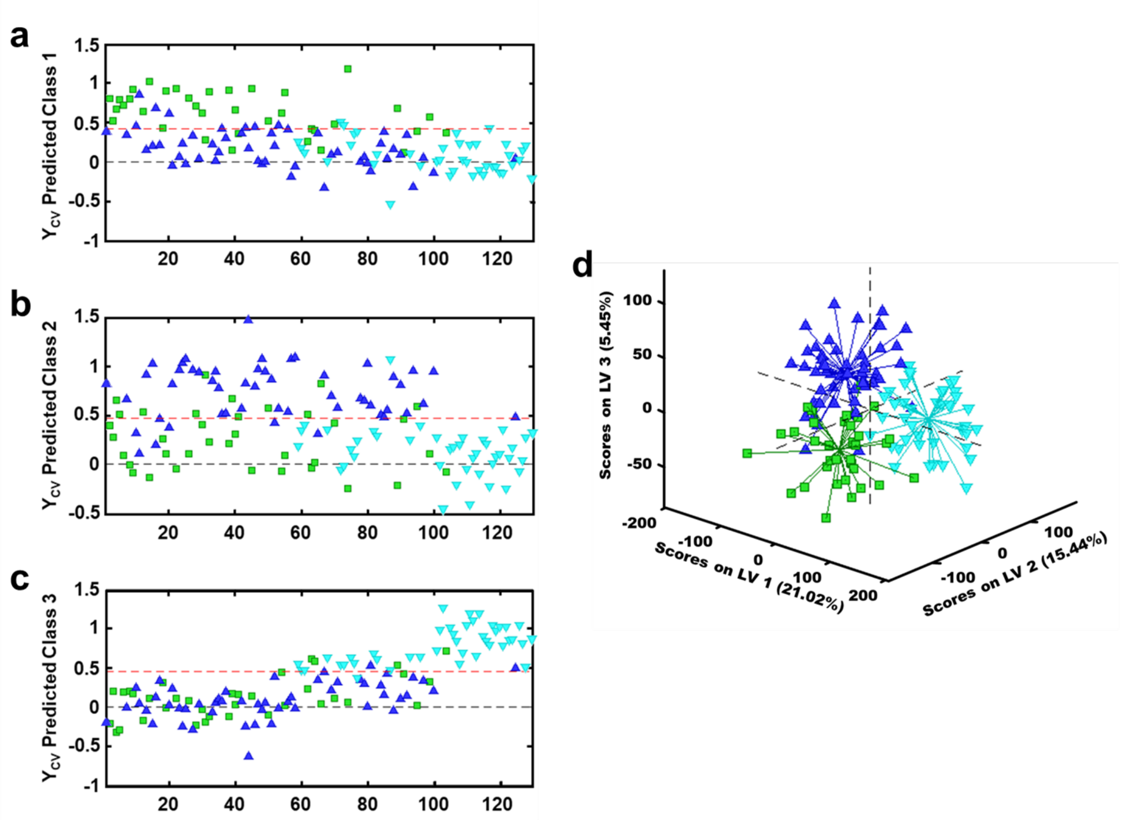

Supplementary Figure 10: Partial Least Squares - Discriminant Analysis (PLS-DA) model using the full timetrace datasets of the rsAKARev, EKARev and RCaMP responses (% increase). (a-c) Show the cross-validation (CV) predictions, where each datapoint (x-axis) is colored according to its classification by the K-means clustering algorithm (green squares, blue triangles, cyan triangles, respectively). The red dotted line indicates the boundary above which a datapoint is correctly classified with 95% confidence. (d) Projection of all datapoints on the 3-LV PLS-DA model. The explained variance (%) of each LV in the original dataspace is shown within the axis labels.

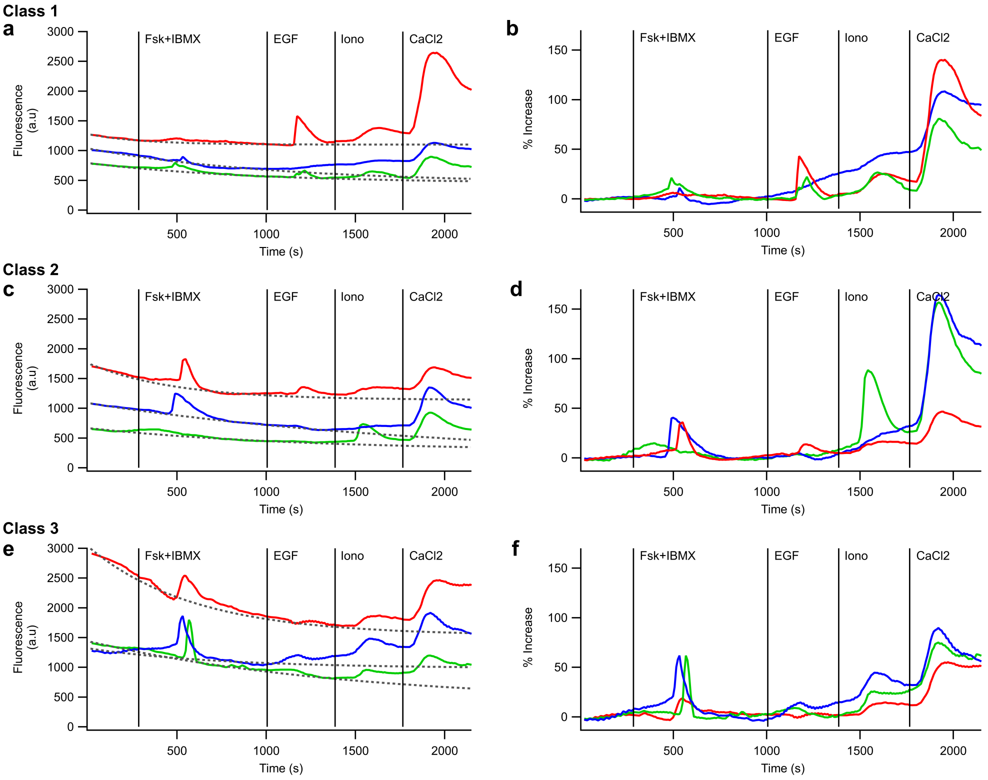

Supplementary Figure 11: Example RCaMP intensity traces, corresponding baselines and calculated responses for representative cells of the different classes. Left panels (a,c,e) show the raw intensity traces (colored, full lines) and corresponding baselines (grey, dashed lines). Right panels (b,d,f) show the quantified responses (% increase) using the methodology described in the methods section. Within the data shown for each class, corresponding data between left and right panels have the same color. Timepoints of pharmacological stimulation are indicated by vertical lines. Cells were stimulated with forskolin (Fsk, 50 µM) and IBMX (100 µM), EGF (1 µg/mL), ionomycin (Iono, 10 µM) and $CaCl_2$ (2 mM).

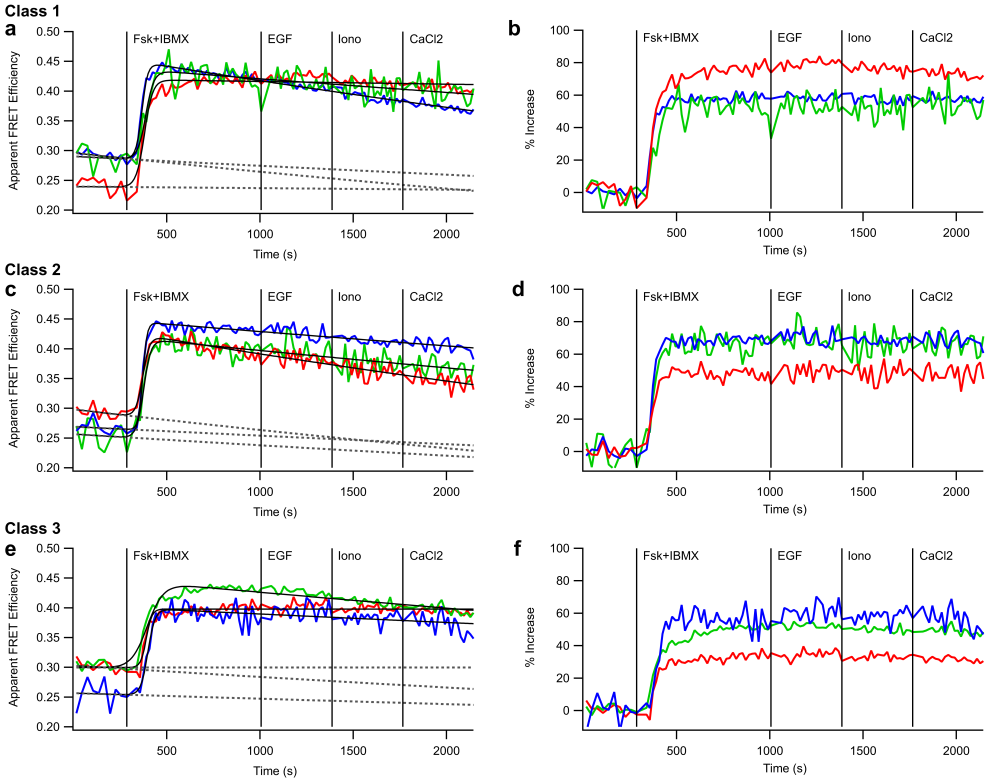

Supplementary Figure 12: Example rsAKARev FRET efficiency traces, corresponding baselines and calculated responses for representative cells of the different classes. Left panels (a,c,e) show the apparent FRET efficiency traces (colored, full lines) and corresponding baselines (grey, dashed lines). Right panels (b,d,f) show the quantified responses (% increase) using the methodology described in the methods section. Within the data shown for each class, corresponding data between left and right panels have the same color. Timepoints of pharmacological stimulation are indicated by vertical lines. Cells were stimulated with forskolin (Fsk, 50 µM) and IBMX (100 µM), EGF (1 µg/mL), ionomycin (Iono, 10 µM) and $CaCl_2$ (2 mM).

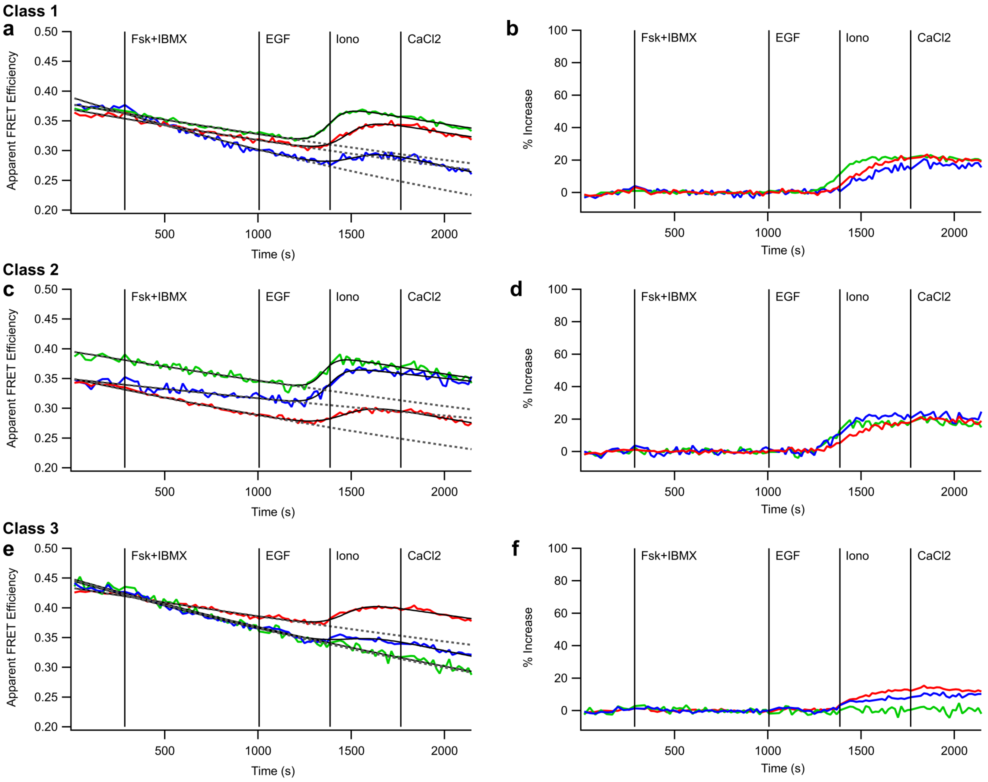

Supplementary Figure 13: Example EKARev FRET efficiency traces, corresponding baselines and calculated responses for representative cells of the different classes. Left panels (a,c,e) show the apparent FRET efficiency traces (colored, full lines) and corresponding baselines (grey, dashed lines). Right panels (b,d,f) show the quantified responses (% increase) using the methodology described in the methods section. Within the data shown for each class, corresponding data between left and right panels have the same color. Timepoints of pharmacological stimulation are indicated by vertical lines. Cells were stimulated with forskolin (Fsk, 50 µM) and IBMX (100 µM), EGF (1 µg/mL), ionomycin (Iono, 10 µM) and $CaCl_2$ (2 mM).
